# Gene-based mapping of trehalose biosynthetic pathway genes reveals association with source- and sink-related yield traits in a spring wheat panel

**DOI:** 10.1101/2020.07.07.192054

**Authors:** Danilo H. Lyra, Cara A. Griffiths, Amy Watson, Ryan Joynson, Gemma Molero, Alina-Andrada Igna, Keywan Hassani-Pak, Matthew P. Reynolds, Anthony Hall, Matthew J. Paul

## Abstract

Trehalose 6-phosphate (T6P) signalling regulates carbon use and allocation and is a target to improve crop yields. However, the specific contributions of trehalose phosphate synthase (TPS) and trehalose phosphate phosphatase (TPP) genes to source- and sink-related traits remain largely unknown. We used exome-capture sequencing on TPS and TPP genes to estimate and partition the genetic variation of yield-related traits in a spring wheat (*Triticum aestivum*) breeding panel with diverse genetic heritage. Twelve phenotypes were directly correlated to TPS and TPP genes including final biomass (source) and spikes and grain numbers and grain filling traits (sink) showing indications of both positive and negative gene selection. Additionally, individual genes explained a substantial proportion of heritability (e.g. 3, 12, and 18% of the variance in gene homeologues most closely related to Arabidopsis *TPS1* for final biomass), indicating a considerable contribution of this regulatory pathway to the phenotypic variation. Most importantly, two significant missense point mutations in the exon 6 of the *TPS1* gene on chromosome 1D substantially increased plant height and peduncle length which was inversely related to grains per m^2^. Gene-based prediction resulted in significant gains of predictive ability (6% improvement) for grain weight when gene effects were combined with the whole genome markers, potentially helping breeding programs in designing strategic crosses. Three *TPS1* homeologues were particularly significant in trait variation. Our study has generated a wealth of information on the role of natural variation of TPS and TPP genes related to yield potential.

## Introduction

In recent decades, numerous studies in bread wheat (*Triticum aestivum* L.) have focused on relations between the source (the supply of assimilates) and the sink (the capacity to accumulate available carbohydrates) with the aim to increase genetic gains (Reynolds *et al*., 2017). A pathway within crops that has been associated with source- and sink-related traits is the trehalose biosynthetic pathway consisting of trehalose phosphate synthase (TPS) and trehalose phosphate phosphatase (TPP) genes (Lunn, 2007; Lunn *et al*., 2014; Paul *et al*., 2018; Paul *et al*., 2020).

The trehalose pathway has a central and indispensable function due to the role of the intermediate trehalose 6-phosphate (T6P) as a signal of sucrose availability (Lunn *et al*., 2006; Schluepmann *et al*., 2003). T6P is an inhibitor of SnRK1 (Zhang *et al*., 2009), a member of the AMPK/SNF1 group of protein kinases that coordinate cellular and organismal responses to carbon and energy. Through SnRK1, T6P de-represses gene expression for carbon use in biosynthetic pathways and growth and development (Nunes *et al*., 2013; Zhang *et al*., 2009). On this basis, the pathway is an ideal candidate for potential modification in crops to alter growth, development, architecture (for yield and yield resilience to abiotic stress), and the biosynthetic pathways that underpin the accumulation of yield-determining end-products such as starch (Paul *et al*., 2018). For example, overexpression of a TPP gene in maize led to improved grain set and yield under a range of water availabilities including mild and severe drought and represents one of few examples of reproducible yield increases in the field through genetic modification of an intrinsic metabolic process in a major crop (Nuccio *et al*., 2015). Kretzschmar *et al*. (2015) showed that a TPP gene, *OsTPP7*, as the genetic determinant in qAG-9-2, a major quantitative trait locus (QTL) for anaerobic germination tolerance. In bread wheat, a TPP gene was associated with grain weight (Zhang *et al*., 2017). Chemical intervention of T6P levels in wheat through the application of UV-cleavable T6P precursors increased grain size in well-watered conditions and enhanced vegetative growth recovery after drought stress (Griffiths *et al*., 2016). All these examples show the centrality of the pathway in determining yield processes. The examples also show the possibility for further optimisation of the trehalose pathway in crops in addition to what may already have been achieved in the development of modern elite varieties through breeding.

New diverse populations of wheat have been developed to incorporate a range of variation from elite, synthetic, and exotic materials aiming at increasing genetic gains associated with a better source and sink balance (Reynolds *et al*., 2007; Reynolds *et al*., 2009a; Reynolds *et al*., 2009b; Reynolds *et al*., 2017). Recently, marker-trait associations have been reported for the High Biomass Association Mapping Panel (HiBAP) consisting of bread spring wheat lines constructed from elite high-yielding material, pre-breeding lines, landraces and synthetically derived lines crossed and selected for high yield and biomass (Molero *et al*., 2019). However, despite some major contributions of association studies, wheat geneticists are increasingly shifting their attention towards the variation inside genes (i.e. exome variants) to look for novel gain-of-function mutations that produce agriculturally useful traits for crop breeding (Gardiner *et al*., 2018; Jordan *et al*., 2015; Uauy *et al*., 2017). Nonetheless, the genetic contribution of specific regulatory pathways to complex traits remains unexplained. Therefore, assessing the cumulative effects of genetic variants in a genic region, partitioning gene heritability, and evaluating the patterns of intragenic linkage disequilibrium (LD) and signatures of selection are important implications to be considered to dissect the genetic architecture of quantitative traits (Timpson *et al*., 2018; Visscher *et al*., 2008). Also, pleiotropy (Visscher and Yang, 2016) and gene-gene interactions (Mackay, 2014) are expected in central regulatory pathways as multiple traits are regulated through the same network(s) (Boyle *et al*., 2017; Hassani-Pak *et al*., 2020).

Gene-based association analysis (Neale and Sham, 2004) is the most common approach to test local regions with increased statistical power compared to single point scans (Lee *et al*., 2014). Several methods have been successfully applied in both simulations and empirical human data (Auer and Lettre, 2015; Bansal *et al*., 2010), but studies in plants are still scarce (Fritsche *et al*., 2012). The most popular approaches used are the sequence kernel association test (SKAT) (Wu *et al*., 2011) and the optimal test named SKAT-O (Lee *et al*., 2012). Nevertheless, there is no clear consensus in the literature about the optimal statistic test once they account for different genetic assumptions (e.g. direction of variant effect) (Bomba *et al*., 2017). Another strategy in quantifying the contribution of a gene region is partitioning the regional genetic variance in combination with the effects of the whole genome (Uemoto *et al*., 2013). Previous studies reported a substantial proportion of heritability captured by breaking down the genome into local bins (Müller *et al*., 2019; Resende *et al*., 2017).

With the advance of exome-capture (enrichment) sequencing (Winfield *et al*., 2012), genome-wide screens for signatures of positive selection have been largely documented in wheat (Russell *et al*., 2016), but selection signals in specific regulatory genes are currently lacking. Besides, traditional implementations of selective pressure analysis (e.g. *d*_N_/*d*_S_ ratio) have typically used simplistic substitution models, potentially leading to increased false-positive rates (Lawrence *et al*., 2013). Therefore, algorithms accounting for variation in sequence context, mutation rates, and trinucleotide mutability have been recently proposed for cancer data sets (Martincorena *et al*., 2017), and application in plants is encouraged. Interestingly, TPS and TPP genes are listed as genes that have been modified during domestication in maize (Hufford *et al*., 2012), potato (Xu *et al*., 2017), and sugarcane (Hu *et al*., 2020). Finally, further useful application of genic variants in applied breeding is the implementation of regulatory pathways in gene-based prediction in maize (Zhang *et al*., 2020). Recent studies applying genomic prediction have demonstrated that predictive abilities can be improved by accounting for gene expression profiles and metabolic abundance information (Dan *et al*., 2016; Schrag *et al*., 2018; Yadav *et al*., 2019). However, the limiting costs associated with those data limited their practical use in crop breeding (Bassi *et al*., 2016). Therefore, a promising strategy is using the contributing genes to predict complex traits in genomic models (Zhang *et al*., 2020).

In this study, we used enrichment-capture sequencing on 25 TPS and 31 TPP genes (Paul *et al*., 2018) for the dissection of the genetic architecture of 24 traits in the wheat HiBAP panel (Molero *et al*., 2019). Our goals were to (*i*) combine single variant analysis and gene-based approaches for the detection of genetic associations, (*ii*) evaluate the intragenic patterns of signatures of selection, (*iii*) estimate the contribution of single gene and the gene families across and within exotic-derived and elite subpopulations, and (*iv*) explore various approaches of gene-based prediction using TPS and TPP kernels.

## Results

### Exome sequences from trehalose pathway genes revealed substantial within-group variation in elite germplasm

We generated the gene ontology network and a framework of the trehalose biosynthetic pathway (Figure 1a). Genome-wide (*F*_*ST*_ = 0.1) and exome profiles (*F*_*ST*_ = 0.06) revealed mild genetic structuring between-groups and a substantial within-group variation in elite lines, suggesting that these genotypes may have some gene-specific mutations (Figures 1b-d). Interestingly, the most contributing alleles to classify the groups were predominantly TPP variants (Figure 1e). The gene-wide LD decayed relatively faster in the complete panel, and regardless of the similar pattern between groups (Figure 1f), elite lines revealed a larger LD block between genes compared to exotic materials, indicating fewer recombination events, mostly due to bottleneck created by the development of elite inbred lines (Figure 1g).

**Figure 1.**
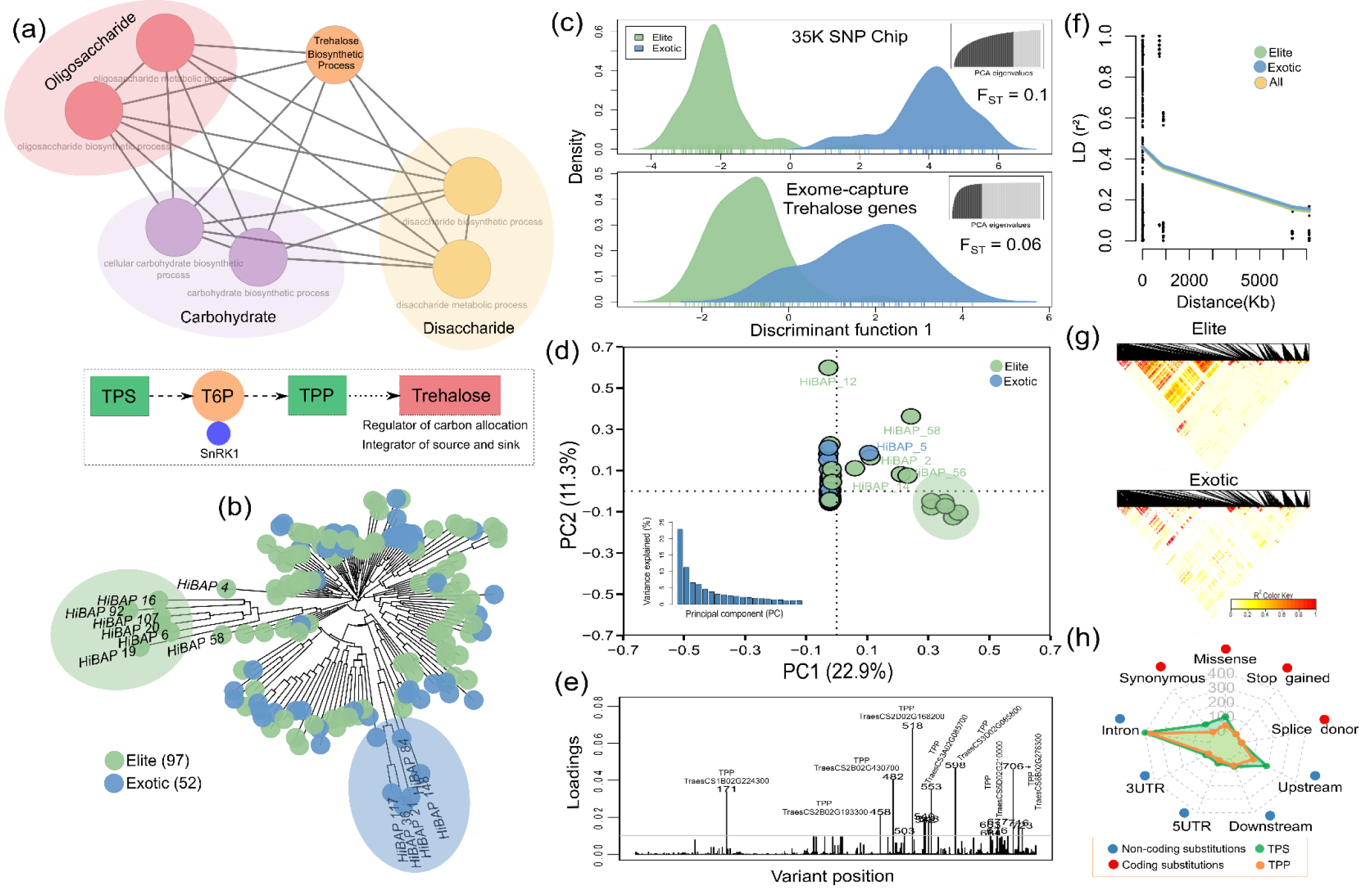
Population structure analysis using the exome-capture data in the wheat HiBAP panel. (a) Gene ontology network and summary of the trehalose biosynthetic pathway. The same color nodes represent similar biological processes. Trehalose phosphate synthase (TPS), trehalose 6-phosphate (T6P), and trehalose phosphate phosphatase (TPP). (b) Neighbor-joining tree (NJT) based on Euclidean distance where each color represents a group (97 elite and 52 exotic derived individuals). Group clustering was determined by Molero *et al*. (2019). (c) Density of individuals from a single discriminant function using the 35K SNP Chip and exome-capture data. Dark grey color on the top right is the number of principal components (PC) retained for the discriminant analysis (DA). *FST* values are shown inside the plots. (d) First two PCs using exome data colored by groups. Bottom left plot represents the variance explained by the first twenty PCs. (e) Contributions (loadings) of each gene variant to the DA function. (f) Pattern of linkage disequilibrium (LD) decay among all pairs of genetic variants for the complete set of individuals (all), elite, and exotic materials. Values reported are the average squared correlations (*r*^2^) across all genes. (g) LD heatmap of the gene variants for elite and exotic subgroups. The color gradient scale represents the range of *r*^2^ values. Black represents the highest estimates of LD. (h) Radar plot showing the distribution of Variant Effect Predictor (VEP) consequences (five non-coding and four coding substitutions) for the trehalose phosphate synthase (TPS) and trehalose phosphate phosphatase (TPP) gene family.

### Gene-based scanning detected multiple trehalose pathway genes associated with key agronomic traits

Exome-capture data showed that the number of variants in TPS and TPP sequences varied greatly among genes from 1-173 in 21 TPS and 27 TPP homeologues (Tables S1-S2), and most of them were predicted as non-coding substitutions (e.g. introns and upstream) whereas, in exonic regions, missense (non-synonymous) substitution was the most prevalent annotation (Figure 1h; Tables S1-S2).

From the single point scans, we detected multiple low-frequency variants in six genes linked with yield-related traits (Table S3). Interestingly, many significant signals identified fell predominantly in non-coding regions, but the most impacting variants were in protein-coding sites. We detected two missense point mutations in the exon 6 of the *TPS1* gene on chromosome 1D that substantially increased plant height and peduncle length but was inversely related to grains per m^2^. One mutation causes a substitution of asparagine to histidine (T → G transversion on site 47313394) while a second threonine to alanine (T → C transition on site 47313412).

The gene-based mapping detected more signals than single variant analysis, identifying a total of 11 TPS and six TPP genes associated with 11 phenotypes (Figure 2; Figure S1). However, some unique associations were only detectable using single point mapping (Table 1). Plant height was the trait with more significant associations (i.e. four TPS and two TPP) followed by peduncle length. There were also effects on grain traits related to spikelet fertility such as number of spikelets per spike, and grains and spikes per m^2^. Despite some variability in the relative performances of the region-based models, we observed a slight advantage of the multiple linear regression approach for gene discovery (Figure 2).

**Table 1.**
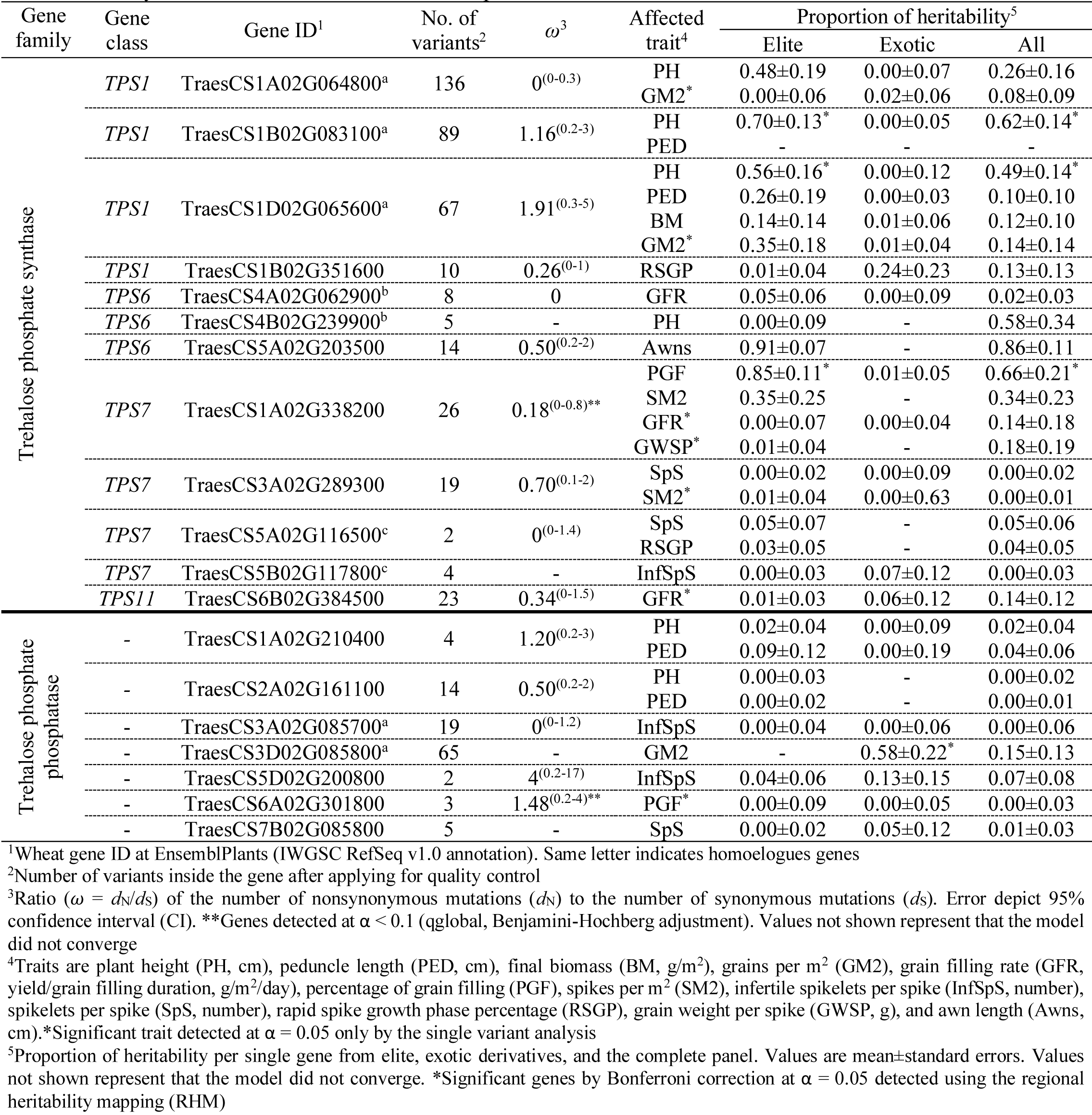
Summary of trehalose phosphate synthase (TPS) and trehalose phosphate phosphatase (TPP) genes significantly associated with yield-related traits in the wheat HiBAP panel

**Figure 2.**
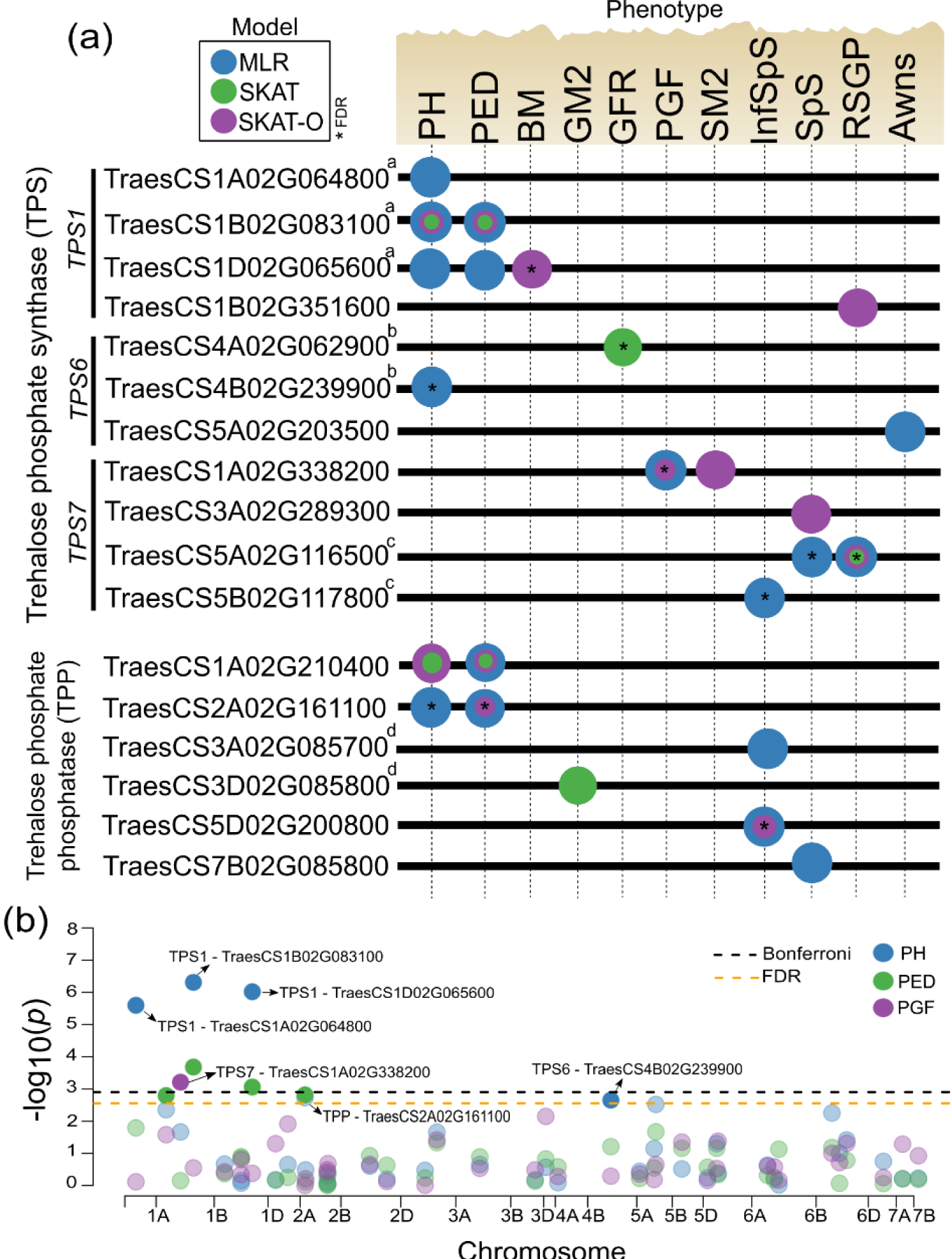
Summary of the gene-based association analysis in the wheat HiBAP panel. (a) Trehalose phosphate synthase (TPS) and trehalose phosphate phosphatase (TPP) genes are shown in the left panel. Same letter indicates homoelogues genes. Circle forms encoded by different colours represent genes detected by sequence kernel association test (SKAT), optimized SKAT (SKAT-O), and multiple linear regression (MLR) models. Significance level used is Bonferroni correction (no asterisk, α=0.05) and False Discovery Rate (with asterisk, α=0.05). Circles overlapping each other represent multiple models detecting the same gene. Only genes and traits on which significant associations were detected are shown in the figure. ^a-d^Same letter right to the gene ID indicates homoelogues genes. (b) Manhattan plot from the gene-based mapping. Results shown are from the MLR model. The *x*-axis shows genomic position (chromosomes 1A-7B), and the *y*-axis shows statistical significance [–log10(*P*)]. Dotted line indicates significance level for Bonferroni correction and False Discovery Rate (α=0.05). Each dot represents a gene coloured by phenotype. Significant gene names are shown by black arrows. Traits are plant height (PH, cm), peduncle length (PED, cm), final biomass (BM, g/m^2^), grains per m^2^ (GM2), grain filling rate (GFR, yield/grain filling duration, g/m^2^/day), percentage of grain filling (PGF), spikes per m^2^ (SM2), infertile spikelets per spike (InfSpS, number), spikelets per spike (SpS, number), rapid spike growth phase percentage (RSGP), and awn length (Awns, cm).

### A significant dominant missense mutation in the *TPS1* gene on chromosome 1D contributed to increasing plant height and peduncle length and was inversely related to grains per m^2^

We classified the wheat panel lines into seven groups with diverse sizes based on allelic combinations of deleterious missense variants on *TPS1* homeologues (Figure 3). On average, the lines (HiBAP 20, 92, 107) carrying the dominant allele on chromosome 1D (group 4) consistently increased plant height and peduncle length compared to those with the alternative allele but reduced the phenotypic values of grains per m^2^ (Figure 3; Table S3). On the other hand, lines (HiBAP 6, 16, 19) carrying only dominant missense mutations on chromosomes 1A and 1B (group 3) showed the opposite behaviour.

**Figure 3.**
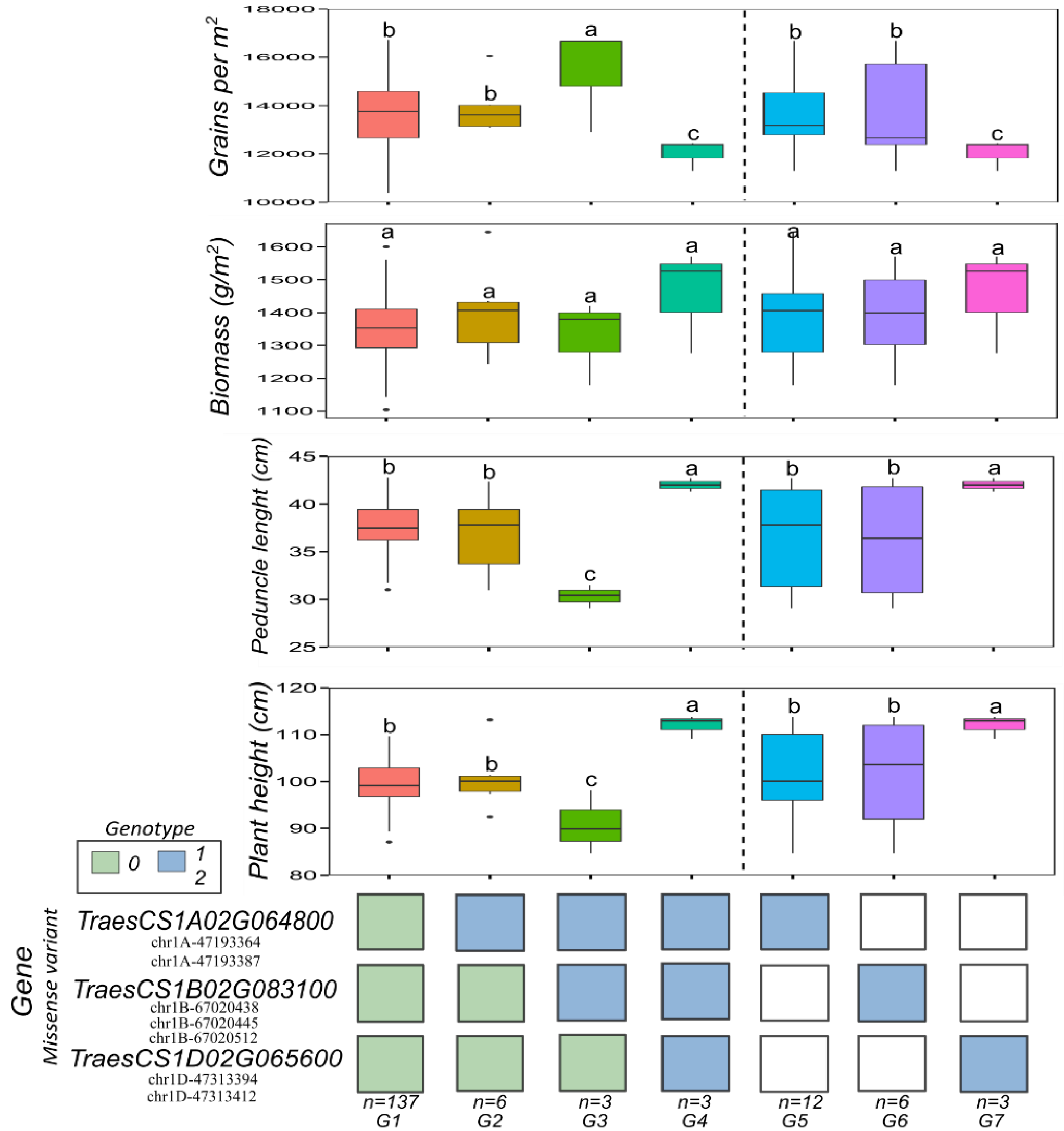
Impact of deleterious missense mutations in the trehalose phosphate synthase (*TPS1*) homoelogue genes to complex traits in the wheat HiBAP panel. Phenotypic differences between individuals carrying different alleles of genetic variants within the gene associated with plant height (PH, cm), peduncle length (PED, cm), final biomass (BM, g/m^2^), and grains per m^2^ (GM2). Blue rectangles represent homozygous (code 2) / heterozygous (code 1) dominant genotypes, and green rectangles represent homozygous (code 0) recessive genotypes. White rectangles represent no defined allele. Only existing allelic combinations are shown. The number of individuals (*n*) for each combination is given under the panel. The missense variant ID is given under each gene name on the bottom left panel. Different letters above box plots indicate significant differences at α = 0.05 from Tukey’s test.

### Trehalose pathway genes revealed positive epistatic interactions, pleiotropy, and distinct intragenic linkage disequilibrium patterns

We identified epistatic interactions within- and between-trehalose pathway genes associated with five yield-related traits, particularly for plant height and peduncle length (Figure 4a, b; Table S1). A large fraction of the interactions was positively associated with the traits (Figure 4c). Accordingly, we found connectivity between coding genetic variants e.g. missense (TPS on chromosome 1A) and synonymous (TPP on chromosome 3D) variants positively interacting with each other to enhance the percentage of grain filling. Furthermore, our results also showed that six genes affected multiple distinct phenotypic traits (i.e. pleiotropic effects), particularly the *TPS1* on chromosome 1D (TraesCS1D02G338200, Figure 2a).

**Figure 4.**
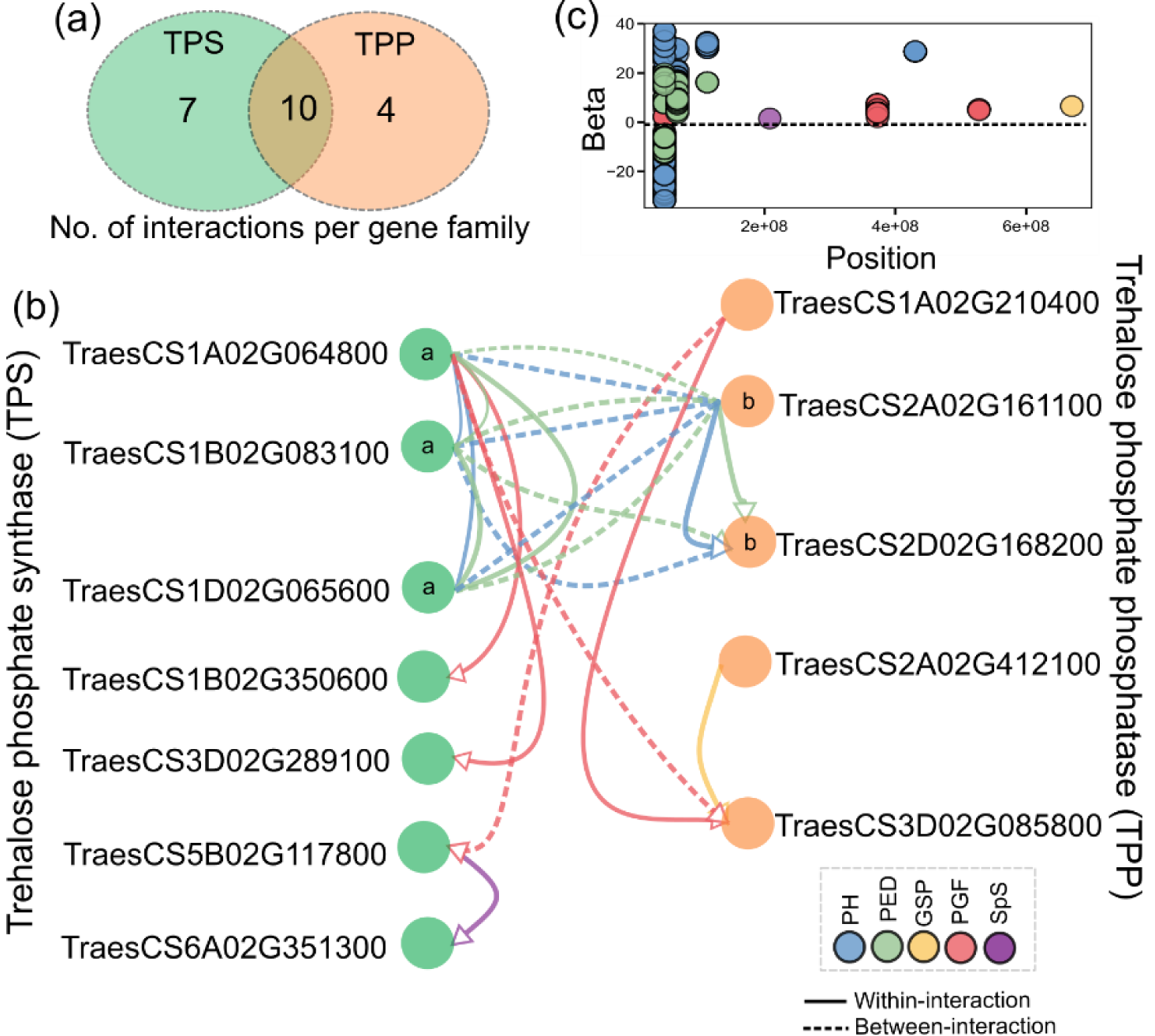
Epistatic interaction of polygenic traits across trehalose family genes in the wheat HiBAP panel. (a) Venn diagram shows the unique and shared number of gene interactions per family. (b) Significant SNP-SNP interaction within-(solid arrow) and between-(dotted arrow) trehalose phosphate synthase (TPS) and trehalose phosphate phosphatase (TPP) gene families. Arrows encoded by different colours represent phenotypes in which interactions between gene variants were found in at least one occasion. Significance level used was Bonferroni correction (α=0.05). ^a-b^Same letter inside circle indicates homoelogues genes. Only genes and traits on which significant connections were detected are shown in the figure. (c) Distribution of regression coefficients (betas) of the interaction against position (bp) of one variant for five phenotypes. Traits are plant height (PH, cm), peduncle length (PED, cm), grains per spike (GSP, number), percentage of grain filling (PGF), and spikelets per spike (SpS, number).

By evaluating the extent of intragenic LD, we identified substantial variation among TPS and TPP genes (Figure S2). For instance, the LD in homeologues followed distinct patterns with some persisting across longer distances (e.g. TraesCS1A02G064800) while others increased with physical distance (TraesCS1B02G083100) or decayed within 1000 base pairs (TraesCS1D02G065600).

### A large fraction of trehalose pathway genes are under positive and negative selection

We measured the strength and mode of natural selection acting on regulatory regions via the *d*_N_/*d*_S_ ratio using a trinucleotide substitution model (Figure 5; Table 1). Nearly one-half of the TPS and TPP genes showed strong indications of negative and positive selection indicating that purifying (negative) and diversifying (positive) selection are acting on them (Figure 5a). We observed high values (*ω*>1, positive selection) of global information (i.e. variation of the mutation rate across genes in each sub-genome) for the chromosomes 1D and 2B (Figure 5d). Interestingly, several genes found to be under positive selection were also associated with a specific phenotype e.g. *TPS1* on chromosomes B and D (plant height, final biomass, and grains per m^2^) (Figure 2; Figure 5) and TPP genes TraesCS6A02G301800 (percentage grain filling) and TraesCS5D02G200800 (infertile spikelets per spike). Some TPP genes showing positive selection were not linked to any traits measured in our study.

**Figure 5.**
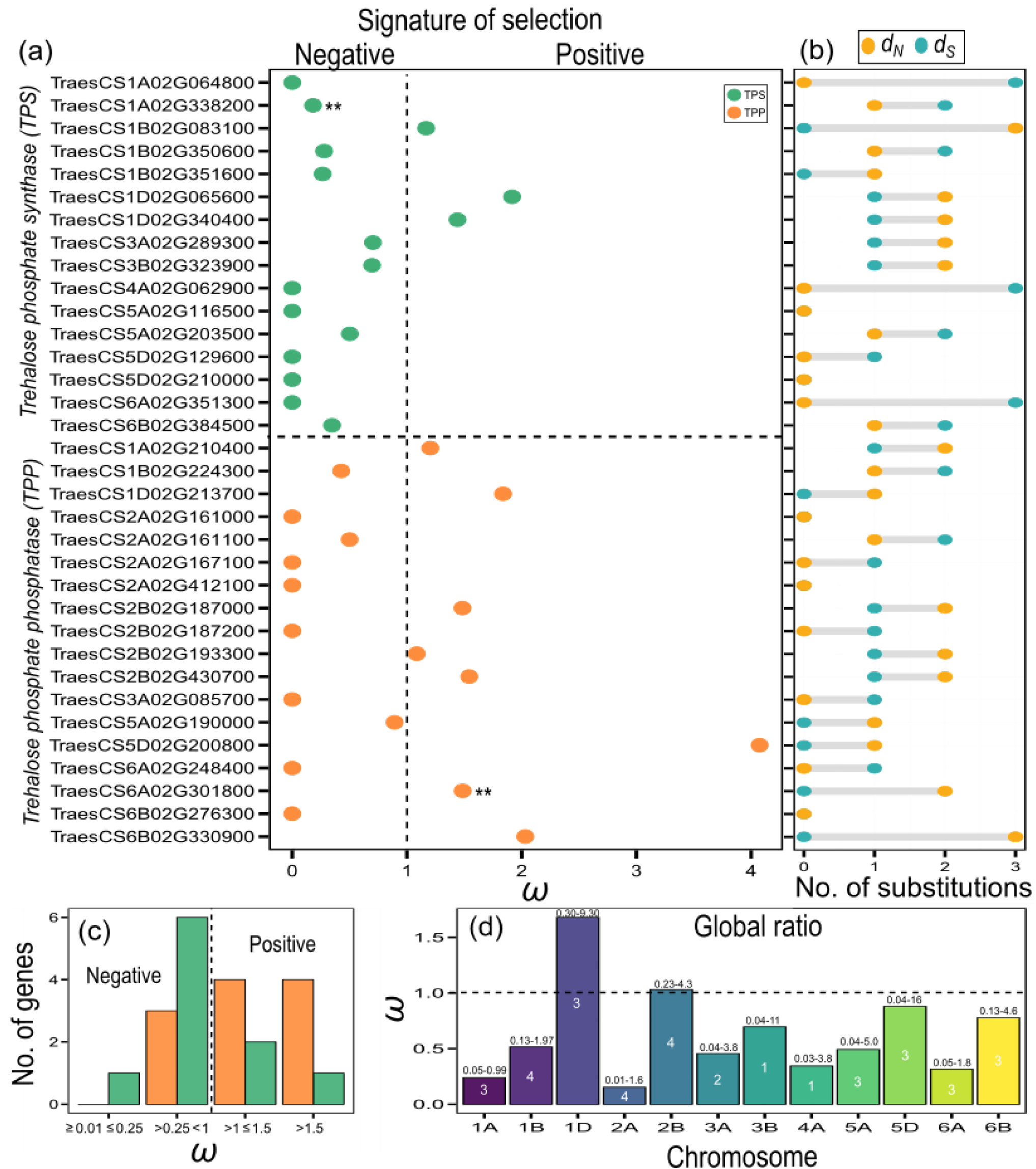
Inference of signature of selection across two trehalose family genes in the wheat HiBAP panel. (a) Gene-wide ratio (*ω*) showing evidence of negative and positive selection. Only genes on which a ratio was estimated are shown in the figure. (b) Total number of non-synonymous (*d*_N_) and synonymous (*d*_S_) substitutions per gene using the *dNdScv* method. **Genes detected at α ≤ 0.1 (qglobal). Trehalose phosphate synthase (TPS) and trehalose phosphate phosphatase (TPP) genes are shown in the panel. (c) Number of genes under different levels of positive and negative selection based on the *ω* ratio distribution. (d) Global *ω* estimates across all genes per chromosome. Values above bar plots indicate 95% confidence interval (CI). The number of genes (*n*) in each sub-genome is given inside the plot.

### Partitioning the total genetic variance of trehalose genes revealed substantial contributions of the pathway to the phenotypic variance

By quantifying the contribution of each variant to the genetic variance of the trait, we observed contrasting patterns of the beta densities, clearly showing differences in the peak and distribution across families (Figure 6a; Table S1). Additionally, most of the variants with large effect sizes were found at low frequencies, following a decay curve shape (Figure 6b, Table S1).

**Figure 6.**
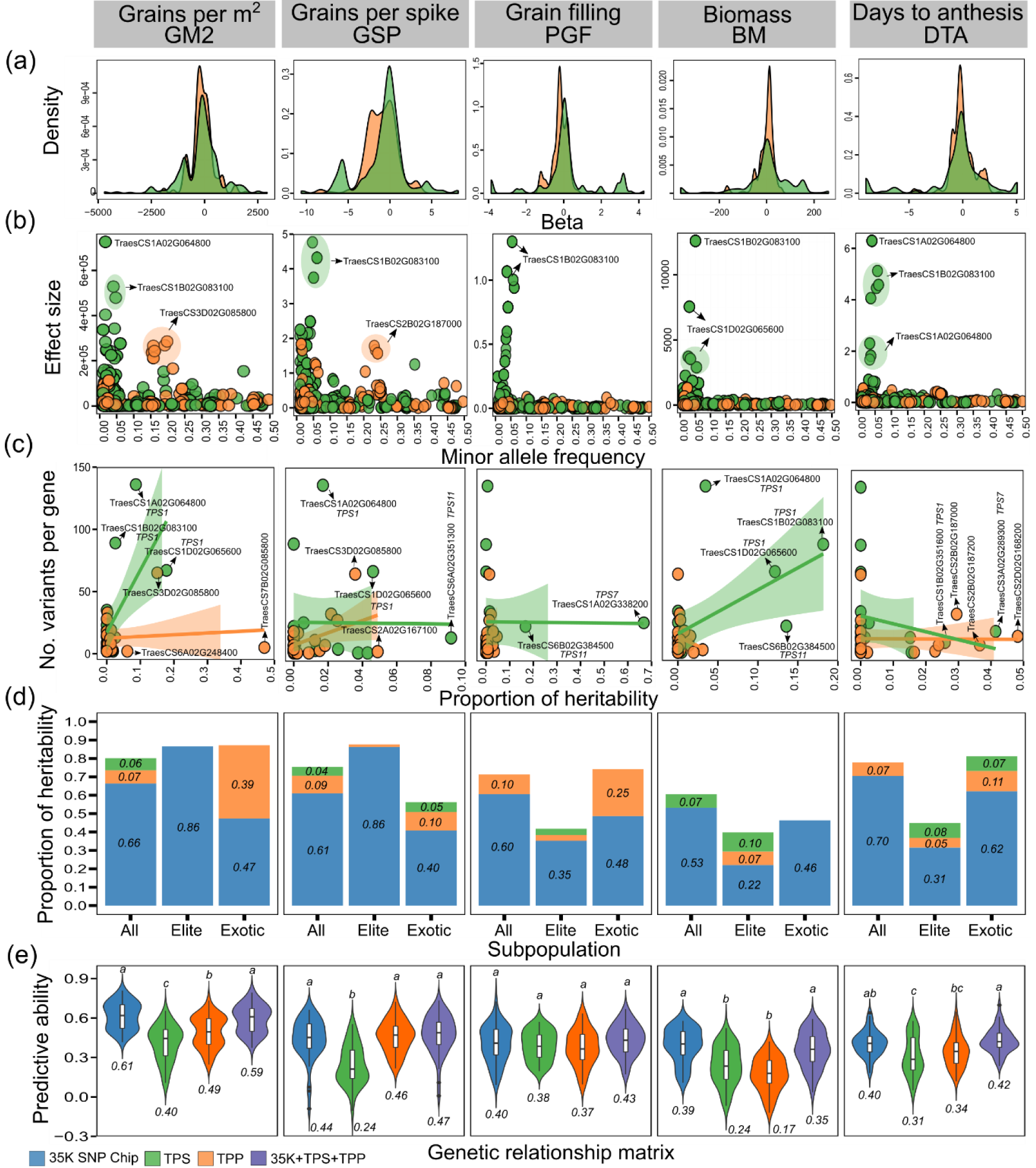
Genetic architecture of complex traits using exome-capture data of trehalose genes in the wheat HiBAP panel. Gene families are trehalose phosphate synthase (TPS) and trehalose phosphate phosphatase (TPP). (a) Estimate of the density distribution of regression coefficients (betas). (b) Association between effect size (ES) and minor allele frequency (MAF). (c) Association between the number of variants and the proportion of heritability per gene. Some genes are shown by black arrows. (d) Proportion of heritability in the complete set, elite, and exotic subgroups. (e) Predictive ability based on genome-wide markers, single gene family, and combined effects. Numbers inside plots represent the mean from 50 random cross-validations. Different letters above violin plots indicate significant differences at α = 0.05 from Tukey’s test. Genome-wide markers represent the 35K Affymetrix Axiom® HD wheat array. Traits are grains per m^2^ (GM2), grains per spike (GSP, number), percentage of grain filling (PGF), final biomass (BM, g/m^2^), and days to anthesis (DTA, days).

We assessed the genetic architecture of complex traits by breaking down the total variance into single gene and gene families, thereby estimating the contribution of each component to the phenotypic variation (Figure 6c, d; Table 1; Table S1). The proportion of heritability per gene varied considerably between gene families (e.g. explaining up to 18% of the variance for biomass) (Figure 6c). Likewise, we showed that *TPS1* homeologues explained a high fraction of heritability (with some significant regions) for the agronomically relevant traits thousand-grain weight, grains per m^2^, plant height, final biomass, and harvest index (Table 1). We further observed a meaningful amount of the variance explained by the TPS family in the complete set (e.g. 0.13 of the heritability for grains per spike) (Figure 6d). Intriguingly, a pronounced contribution (e.g. 0.02-0.41 of the heritability) of TPP genes was evidenced in the exotic germplasm, particularly for sink traits e.g. grains per m^2^, grains per spike and percentage of grain filling.

Under the expectation that well-known regulatory pathways could be used in gene-based prediction, we identified that complex phenotypes were moderately predicted using only single-family effects (e.g. 0.09-0.47 for TPS, and 0.03-0.49 for TPP) (Figure 6e; Table S1). When we included gene effects simultaneously with whole-genome markers we observed relatively little gains of predictive ability, except for grain weight per spike (i.e. significant increase of 6% compared to the traditional model).

## Discussion

The T6P signalling pathway (Figure 1a) is a central regulatory system of resource allocation and source-sink interactions and is emerging as an important target in crops such as maize, rice, wheat, and sorghum (Paul *et al*., 2018; Paul *et al*., 2020). Here for the first time we analysed comprehensive exome SNP information for TPS and TPP genes and dissected the genetic architecture of yield-related traits in a spring wheat diversity panel where all lines presented acceptable agronomic performance (Molero *et al*., 2019). The data showed significant relationships of TPS and TPP genes with agronomic traits with evidence of historical selection and identified opportunities for future selection of TPS and TPP genes and strategic crossing for yield improvement.

### Gene-based scanning detected multiple trehalose pathway genes associated with key agronomic traits

Previous genome-wide and exome studies of complex traits suggested that both coding and non-coding variants tend to contribute to the phenotypic variance (Li *et al*., 2016; Visscher *et al*., 2017). Consistent with this expectation, we identified both substitution types (mostly intronic regions) associated with the phenotypes (Table S3). Moreover, we empirically confirmed the argument that gene-based mapping would have greater statistical power than conventional single analysis (Li and Leal, 2008). However, on a few occasions, the latter resulted in the detection of additional signals, presumably because of the limitations associated with the region-based testing (e.g. differences in the local allelic architecture of a trait) (Timpson *et al*., 2018). We therefore recommend using both analyses to maximize the overall number of variants/genes detected. We also observed that the multiple linear regression approach often outperformed the methods with random-effects (Figure 2a) showing relatively good model fit (see QQ plots in Figure S1) although false positives are often reported for fixed-effect models (Deng and Pan, 2018; Svishcheva *et al*., 2019).

We did not detect any signals coming from the trehalose biosynthetic pathway in the association mapping using the same HiBAP panel where exome-capture sequencing was not utilised (Molero *et al*., 2019). Therefore, exomic data is a viable option of narrowing down the relative importance of core network genes (Kiezun *et al*., 2012) as most of the GWAS hits are outside of genic regions (Visscher *et al*., 2017).

### A significant dominant missense mutation in *TPS1* gene on chromosome 1D contributed to increasing plant height and peduncle length and was inversely related to grains per m^2^

We found two deleterious missense variants leading to amino acid changes in the *TPS1* gene on chromosome 1D (Table 1; Figure 3). This finding likely suggests that these mutations in a single homeologue led to a gain-of-function allele that effectively dominates the other two homeologues (Borrill *et al*., 2019). Further studies are required to confirm this (Zhu and Qian, 2020) and to understand how the mutation affects the function of *TPS1*, through its expression or catalytic properties. Strong effects on phenotype have been shown in *Arabidopsis thaliana* where mutants carrying weak alleles of *AtTPS1* showed less T6P than wild-type plants, consequently, reducing vegetative growth (Gómez *et al*., 2010). It is not yet known whether the wheat TPS homeologues closest to *A. thaliana TPS1* perform the same function as *A. thaliana TPS1* in synthesising T6P. Further work would be needed to confirm this.

Despite the general increase in the phenotypic values of height and peduncle length, we also found a consistent reduction of grains per m^2^. This result was expected as the traits are negatively correlated (Molero *et al*., 2019) and regulated through the same causative gene (Table 1). This finding suggests that mutations in the *TPS1* gene have a large effect on whole plant carbon allocation and growth, potentially through SnRK1 which regulates many genes involved in metabolism and resource allocation (Zhang *et al*., 2009). *TPS1* homeologues could be a target for gene editing, particularly if this were able to increase grain numbers.

### Trehalose biosynthetic genes revealed positive epistatic interactions, pleiotropy, and distinct intragenic linkage disequilibrium pattern

Under the assumption that epistasis is relatively common in central genetic networks of highly polygenic traits (Mackay, 2014) we provided evidence of connectivity between variants (mutations) from trehalose pathway genes, particularly for a set of complementary genes on the A, B, and D genomes (Figure 4). For instance, we observed that the *TPS1* homeologues interacted considerably with TPP homeologues on chromosomes 2A and 2D. Similar patterns of trehalose pathway gene interactions have been reported in other organisms e.g. nematodes and yeast (Apweiler *et al*., 2012; Kormish and McGhee, 2005). Likewise, our results also revealed strong pleiotropic effects (Figure 2; Table 1). This can likely be explained by changes in whole plant carbon allocation which may affect more than one trait, particularly for pathways like trehalose that regulates carbohydrate allocation (Paul *et al*., 2018; Paul *et al*., 2020). Previous studies have reported pleiotropy for other phenotypes (e.g. large changes in vegetative architecture and relationships with sucrose content) in trehalose pathway transgenics (Goddijn and van Dun, 1999; Lunn *et al*., 2014; Romero *et al*., 1997).

Contrasting patterns of gene LD have been reported across a range of studies in maize (Ching *et al*., 2002; Remington *et al*., 2001), barley (Caldwell *et al*., 2006), rice (Mather *et al*., 2007), and rye (Li *et al*., 2011), but investigation of wheat genes remain largely unexplored (Sela *et al*., 2011). Thus, by evaluating a large set of exome regions (Figure S2) we identified high levels of intragenic LD (persisting across longer distances), possibly as a result of the reduced recombination rates due to strong artificial selection (Palaisa *et al*., 2003; Remington *et al*., 2001). We also observed a few occasions where LD decayed rapidly within distance reflecting the impact of local recombination. Surprisingly, LD values increased slightly with genomic distance in some genes (due to the large proportion of LD). Similar observations were reported in earlier studies and are likely to be justified by the small effective population size and reduced number of markers per gene (Li and Merilä, 2010; Yang *et al*., 2014).

### A large fraction of trehalose pathway genes are under positive and negative selection

In the screens for signatures of selection, we identified similar proportion of genes under positive and negative selection (Figure 5). Two *TPS1* homeologues showed evidence of positive selection indicating they underwent breeding selection. If it is confirmed that these *TPS1*s regulate T6P levels as catalytically active enzymes, this is consistent with previous observations where alteration of T6P levels through *TPS1* have large effects on phenotype (Avonce *et al*., 2004; Blázquez *et al*., 1998; Wahl *et al*., 2013). Exactly how the wheat *TPS1*s regulate phenotype awaits further work. Additionally, two *TPS7* genes on chromosomes 3A and 3B that are under negative selection in our study were identified to be associated with domestication improvement in the closest genes in maize (Hufford *et al*., 2012; Paul *et al*., 2018).

Interestingly, a large proportion of the positive selection was attributed to the TPP family, indicating that most of the non-synonymous substitutions in these genes might be essentially driver mutations i.e. providing a selective advantage (Pon and Marra, 2015). TPP genes showed positive selection for traits such as percentage of grain filling (Figure 5). On the other hand, traits such grains per m^2^, percentage grain filling, grain filling rate, spikelets per spike, grains per spike and final biomass showed negative selection on many TPS genes, meaning that most of the mutations were removed by negative selection throughout crop breeding (Casillas and Barbadilla, 2017; Vitti *et al*., 2013). For instance, a significant ratio of *ω*∼0.18 indicates that at least ∼82% of missense mutations have been removed by negative selection. Despite clear evidence of genes under selective pressure, only a small fraction showed significant ratios after correcting for multiple testing. Possibly, the limited numbers of mutations per gene lead to noisy estimates of *d*_N_/*d*_S_ values, reducing the power of signature detection (Martincorena *et al*., 2017).

### Trehalose pathway genes revealed substantial contributions to the quantitative genetic variation of source- and sink-related traits

We found that each trehalose pathway gene revealed some proportion of heritability for at least one specific trait (Figure 6c; Table S1), suggesting a substantial contribution of this regulatory biosynthetic pathway to the phenotypic variance. Comparing this result to those from the gene-based testing (Figure 2), we found that the latter yielded fewer associations (e.g. the *TPS1* gene on chromosome 1D is linked to three traits, but explained some variance to twelve traits), presumably because exome signals tend to be difficult to detect with small effect sizes (Sham and Purcell, 2014). Additionally, a *TPS1* gene known to regulate flowering time in Arabidopsis (Wahl *et al*., 2013) explained around 2% of the heritability in our panel (Figure 6c). In a recent study, a TPP gene on chromosome 6A was revealed to be associated with thousand grain weight in bread wheat and successfully cloned (Zhang *et al*., 2017). Similarly, we observed about 3% of heritability fraction in the same TPP gene being associated with this particular trait.

Interestingly, we found heritability per gene consistently higher than expected on a few occasions (roughly explaining 30-86% of the genetic variance) even combined with the whole-genome marker effects (Figure 6c; Table S1). This result was unexpected as a single gene is unlikely to explain more than one-half of the total phenotypic variance, because of the nature of the genetic architecture of polygenic traits (Boyle *et al*., 2017). One main reason for this could be due to our small sample size and/or low coverage of variants per gene, consequently, resulting in overfitting (Evans *et al*., 2018; Yang *et al*., 2011b). Similar patterns of large fraction of additive variance in genomic regions have been reported for complex traits in common bean (Resende *et al*., 2018). Additionally, contrary to some studies (Browning and Browning, 2011; Goddard *et al*., 2011), we observed inflated estimates (in most cases upward) of heritability when accounting for the effect of population structure (Table S1). We further partitioned the gene variance within subgroups, and despite reducing the sample size even more, we found that the *TPS1* homeologues explained a high fraction of heritability in elite materials, particularly for plant height (Table 1).

We also estimated the relative contribution of each gene family to the phenotypic variance of subpopulations (Figure 6d; Table S1). As expected from the quantitative genetics’ theory (Barton *et al*., 2017) a larger proportion of heritability was captured by genome-wide markers. Additionally, we observed contrasting contributions of the trehalose pathway family to elite and exotic materials. Firstly, both TPS and TPP gene families showed very little contributions to exotic germplasm for biomass, plant height, and harvest index, but their proportion increased considerably in elite materials (up to 20% higher), suggesting that the selection process impacted carbon allocation and source and sink balance controlled by trehalose pathway (Paul *et al*., 2018). On the other hand, we observed that the TPP family explained a high fraction of heritability in exotic materials for grain-related traits (Figure 6; Table S1), and not so much for elite lines suggesting potential for inclusion of genetic variation for these traits from exotic germplasm into breeding crosses to increase yield via these traits. Secondly, both gene families showed a high contribution to elite and exotic subpopulations for days to anthesis, suggesting that trehalose genes had little impact on flowering during the selection process in our panel, but suggest the pathway as a whole contributes substantially to flowering time, which was fixed early on in the breeding process. Similar patterns of genetic variation changes through selection have been reported across a range of complex traits (Briggs and Goldman, 2006; Raquin *et al*., 2008).

Our findings on predicting complex traits using their contributing genes have several important implications for designing strategic crosses aiming to improve source-sink balance in wheat. As expected, predicting phenotypes by using single-gene families showed low values of predictive ability, but surprisingly, clearly captured enough information to represent the kinship. Contrasting results have been reported showing a clear advantage of gene-based prediction compared to genome-wide markers (Zhang *et al*., 2020). Our second strategy was combining whole-genome marker effects with causative gene variants, and we observed that predictive ability did not improve significantly in any of the traits, except for grain weight per spike (Table S1). On one hand, genome-wide marker effects are most likely capturing the variation from other core genes, and possibly the information of both types of kinship would be redundant, consequently not effectively contributing to improving prediction (Lyra *et al*., 2019). On the other hand, incorporating gene effects considerably increased predictive ability for grain weight, suggesting that the associated regulatory pathway highly impacted the grain weight phenotype (see the proportion of heritability per single gene and subpopulation), thus adding extra information to the model.

### Potential implications of using the trehalose pathway gene in wheat strategic crossing

We demonstrated the impact of the trehalose biosynthetic pathway to the quantitative genetic variation of yield-related traits allowing the prioritisation of target genes (*TPS1, TPS7, TPS11* homeologues and several TPPs). Selection of genes within the pathway appears to be ongoing and positive for *TPS1* genes and several TPP genes and to have already had a significant impact on harvest index, final biomass, plant height, and flowering time.

Therefore, our findings provide strong evidence that the trehalose gene family will support designing strategic crossing and pre-breeding in various ways. First, we identified several promising genes that in conjunction with gene-editing techniques could be used to study their role in source/sink pathways, e.g. *TPS1* homeologues to elucidate their mode of action within the T6P pathway mechanism. Second, TPP genes in exotic derived material could be reintroduced to enhance grain-related traits e.g. grains per m^2^. Third, there were strong epistatic interactions between genes that could enable gene combinations to be considered e.g for *TPS1* and TPP genes for percentage of grain filling. Fourth, predicting wheat phenotypes by combining whole-genome marker effects with trehalose pathway gene effects has the potential to be a viable predictive model, helping breeding programs to design strategic crosses.

Our study underlines the importance of the trehalose pathway as a contributor to crop improvement both historically and for the future supported here for the first time through comprehensive analysis of diverse genetic variation and its association to yield traits conducted under high yielding conditions. A better understanding of the mechanistic basis of yield improvements achieved by TPS and TPP genes would enable further refinement and advancement of selection and improvement strategies and identify key genes downstream of T6P signalling. Production of near-isogenic lines particularly for interesting genes such as *TPS1* and use ofT6P precursors in chemical genetics (Griffiths *et al*., 2016) could be used to interrogate genetic variation and gene heritability. Further genetic associations will likely be found where traits are determined under different environmental conditions such as abiotic stress.

### Experimental procedures

#### Enrichment capture sequencing and variant calling

Enrichment capture and bioinformatics analysis were carried out as per Joynson *et al*. (2020). Briefly, DNA was extracted from flag leaf material from each panel member. A combined 10 leaves per plot were pooled prior to extraction using a standard CTAB method.

Sequences for 25 TPS and 31 TPP genes were taken from Paul *et al*. (2018). For TPSs these were annotated as *TPS1, TPS6, TPS7*, and *TPS11* in accordance with the nearest *A. thaliana* TPS (Paul *et al*., 2018). For TPPs this was not possible due to greater genetic divergence in TPPs between the two species. Probe sequences (120bp) were designed in an end-to-end format targeting gene bodies and 2000bp upstream ensuring capture of each gene’s promoter sequence. These were integrated into a 12Mb capture probe set. Libraries were constructed using the TruSeq DNA library preparation kit (Illumina) and were sequenced on a NovaSeq6000. The 150bp paired-end sequences were mapped to the Refseq-v1.0 reference sequence using BWA MEM version 0.7.13 73 with subsequent filtering carried out using SAMtools v1.4 and Picard Tools MarkDUplicates. Variants were called using bcftools and filtered using GATK (McKenna *et al*., 2010).

#### Mapping population and phenotypic traits

Mapping population and phenotypic data analyses have been described by Molero *et al*. (2019). Briefly, we used the HiBAP population comprising 149 wheat spring genotypes of the CIMMYT wheat pre-breeding and breeding programme. This panel was divided into two main subpopulations consisting of 97 elite lines and 52 exotic derivatives (landraces, synthetic, and introgression lines). Both groups were selected based on good agronomic performance and high biomass expressed at different times during the growing cycle. The field trials were conducted in two consecutive growing seasons (2015/16 and 2016/17) under fully irrigated conditions situated in the Yaqui Valley, Mexico.

We used the means adjusted for spatial and temporal factors of 24 phenotypes: plant height (PH, cm), peduncle length (PED, cm), biomass at physiological maturity (BM, g/m^2^), harvest index (HI), yield (Yield, g/m^2^), thousand grain weight (TGW, g), grains per m^2^ (GM2), percentage of grain filling (PGF), grain filling rate (GFR, yield/grain filling duration, g/m^2^/day), spikes per m^2^ (SM2), grains per spike (GSP), grain weight per spike (GWSP, g), spikelets per spike (SpS, number), infertile spikelets per spike (InfSpS), spike length (Spike, cm), awn length (Awns, cm), rapid spike growth phase (RSGP, percentage), days to initiation of booting (DTInB), days to anthesis (DTA), days to maturity (DTM), thermal time to initiation of booting (TTInB), thermal time to anthesis (TTAnth), thermal time to maturity (TTPM), and thermal time to anthesis +7days (TTA7H). For further detail on trait evaluation see Molero *et al*. (2019).

#### Variant filtering and annotation

21 TPS and 27 TPP genes showed at least one variant and these were submitted to variant filtering (Table S2). Markers with low minor allele frequency (MAF, <1%) and high call rate (CR, <95%) were removed, and the remaining missing variants were imputed using Beagle 4.1 (Browning and Browning, 2016) within the *codeGeno* function from the Synbreed R package (Wimmer *et al*., 2012). Heterozygous genotypes were kept in the marker data (Allen *et al*., 2013). The final genotypic matrix was composed of 749 variants. The Ensembl Plants (Bolser *et al*., 2016) variant effect predictor (VEP) tool (McLaren *et al*., 2016) was used to annotate variants (coding and non-coding substitutions) and retrieve the functional impact scores of non-synonymous mutations according to the Sorting Intolerant From Tolerant (SIFT) algorithm (Vaser *et al*., 2016).

#### Inference of population structure and genetic differentiation

We explored the gene ontology network (biological process) of the trehalose biosynthetic pathway using the Cytoscape ClueGo plug-in (Bindea *et al*., 2009) inputting the wheat reference genome (IWGSC RefSeq v1.0 annotation) from Ensembl Plants. Additionally, we estimated the genetic diversity of exome variants by calculating the polymorphic information content (PIC) and MAF using the *popgen* function from snpReady R package (Granato *et al*., 2018).

We detected the genomic diversity structure of the population at the gene level. First, we applied a principal component (PC) analysis using the SNPRelate R package (*snpgdsPCA* function) (Zheng *et al*., 2012). Second, we applied a discriminant analysis of principal components (DAPCs) using the adegenet R package (Jombart *et al*., 2010). The group clustering used was inferred by Molero *et al*. (2019). The contributions (loadings) of each gene variant were estimated using the *loadingplot* function. Finally, a neighbor-joining tree (NJT) was generated based on the modified Euclidean distance using the ape R package (Paradis *et al*., 2004) and the pairwise genetic distance between populations (*F*_*ST*_) was calculated following Weir and Cockerham (1984) in the SNPRelate R package. The genome-wide marker data (9267 variants remaining after quality control), generated using the 35K Affymetrix Axiom® HD wheat SNP array (Allen *et al*., 2017), was used only for the DAPCs.

We estimated the level of linkage disequilibrium (LD) between and within trehalose genes using the square allele frequency correlation coefficient (*r*^*2*^) calculated for each pairwise combination in PLINK v.1.9 (Purcell *et al*., 2007). The LD decay curve was fitted by a non-linear regression model (Marroni *et al*., 2011), obtained by fitting *r*^*2*^ with distance using Hill and Weir expectation of *r*^*2*^ between adjacent sites (Hill and Weir, 1988; Remington *et al*., 2001). Haplotype LD block was visualized in elite and exotic subgroups separately using the LDheatmap R package (Shin *et al*., 2006). Pairwise variant interactions between gene regions were tested by a linear regression analysis using PLINK --epistasis command. Regression coefficients (betas) were estimated for each interaction. The Bonferroni multiple testing was used to correct the epistatic significance threshold (0.05/N), where N is the number of interactions tested.

#### Single point scan and gene-based mapping

Single variant association analysis was performed using a Mixed Linear Model (MLM) in GAPIT v3.0 R package (Lipka *et al*., 2012; Wang and Zhang, 2018) incorporating genomic kinship (K) matrix and the first three PCs (Q) to control for the confounding effects of cryptic relatedness and population structure (Yu *et al*., 2006). The default false discovery rate (FDR) (Benjamini and Hochberg, 1995) and Bonferroni multiple testing (Hochberg, 1988) was used to correct the genome-wide significance thresholds (α=0.05).

Following recommendations that region-based tests have different assumptions about the genetic effects and weighting functions (Bomba *et al*., 2017; Lee *et al*., 2014; Nicolae, 2016), we measured the performance of gene mapping empirically using three approaches. First, we used a traditional multiple linear regression (MLR) model (Chapman and Whittaker, 2008) considering genotype effects as fixed. Second, we applied the SKAT model (Chen *et al*., 2013; Wu *et al*., 2011) assigning an Identity by State (IBS) kernel function. Third, we used the combination of burden test and SKAT named SKAT-O (Lee *et al*., 2014; Lee *et al*., 2012). Both kernel-based tests consider the genotype effects as random. Variance components were estimated using restricted maximum likelihood (REML). The weights were calculated using the standard probability density function of the beta distribution. For further detail on the model description see Svishcheva *et al*. (2019). We estimated the *P* values by using Kuonen’s method (Kuonen, 1999) and considered the mode of inheritance as additive. The genomic relationship matrix (GRM) was calculated using the first formula proposed by VanRaden (2008), and the first three PCs were used as covariates in the models. Gene-based mapping was performed using the *MLR* and *FFBSKAT* (*rho* was assigned for SKAT-O test) functions in the FREGAT R package (Belonogova *et al*., 2016). Genes containing only one variant were removed from the analyses. We included all variant annotations (coding and non-coding) in the tests (Neale and Sham, 2004) following suggestions that combining signals from multiple mutations in the same gene increases model statistical power (Sham and Purcell, 2014). We adjusted the *P*-values for multiple comparisons to control for type I error at α = 0.05 using the traditional FDR and Bonferroni procedure (the number of genes tested was considered to set the threshold) using the *p.adjust* R function. Finally, quantile-quantile (Q-Q) plots were used to verify the fitness of the model and plotted using the CMplot R package (https://github.com/YinLiLin/R-CMplot).

We explored allelic combinations for further designed breeding schemes and evaluated the impact of non-synonymous mutations on phenotypic traits. For this, we classified our wheat panel into seven possible groups (G1-G7) based on the genotypes at deleterious missense variants in the trehalose phosphate synthase1 (*TPS1*) homoelogues.

#### Screening for signature of selection at the gene level

We evaluated the evidence of selection at the gene level by estimating the normalized ratio of non-synonymous (missense, nonsense, and splicing) substitutions per synonymous site (*ω = d*_N_/*d*_S_) using an optimized Poisson-based model (*dNdScv*) in the dndscv R package (Martincorena *et al*., 2017). Briefly, this model accounts for variation in mutation rates, sequence context, and full trinucleotide mutability. To estimate the mutation rate of a gene it uses a joint likelihood function combining local (synonymous substitutions in a gene) and global (negative binomial regression across genes) information to estimate the mutation rate of a gene. We used the *buildref* function to input the wheat reference genome (IWGSC RefSeq v1.0 annotation) from Ensembl Plants per chromosome. Global *ω* estimates across all genes were estimated per chromosome. A global *q*-value ≤ 0.1 (without considering InDels) was used to identify statistically significant genes. A confidence interval (α = 0.95) was calculated per gene. Selection was measured as positive (*ω* > 1), negative (*ω* <1), and neutral (*ω* =1) (Nielsen, 2005).

#### Partitioning the genic heritability and predictive models

We investigated distributions of population genetic parameters by estimating beta and effect size only for common variants. First, we estimated the coefficient of regression (β) by fitting a single-point association test (Q+K model) using the FREGAT R package. Briefly, beta is the absolute additive effect of the minor alleles on the phenotype in standard deviations (Park *et al*., 2011; Timpson *et al*., 2018). Second, we estimated the effect size, defined as the contribution of the variant to the genetic variance of the trait, following the equation: *EF* = 2*β*^2^*f*(1 − *f*), where *β* measures the regression effect, and *f* denotes the minor allele frequency (Park *et al*., 2011; Park *et al*., 2010).

We further investigated the genetic architecture of complex traits by partitioning the genetic variation of individual genes and gene families within and across elite and exotic subpopulations using the genomic-relatedness-based restricted maximum-likelihood (GREML) approach (Yang *et al*., 2010) implemented in GCTA software v1.93.1beta (Yang *et al*., 2011a). To estimate the proportion of the phenotypic variance explained (i.e. genomic heritability) per gene we fitted multiple GRM in the model, one contributed by the whole genome (35K SNP Chip) and a second by a specific gene region. The proportion of heritability was estimated ignoring population structure (Table 1; Figure 6) and adjusting PCs as fixed covariates (Table S1). We reported the single gene heritability as 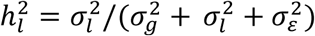 (see Methods S1). Additionally, we partitioned the variation of gene family as 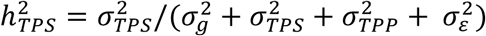 (see Methods S2). 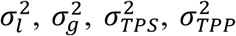, and 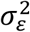 are the local gene, global whole genomic, TPS and TPP gene family, and residual variances, respectively.

We used the additive genomic best linear unbiased prediction (GBLUP) model controlling for population structure (Lyra *et al*., 2018) to compare the predictive ability of four gene-based approaches (see Methods S3). Prediction of the phenotypes was performed by using the (*i*) genome-wide marker (35K SNP Chip) effects, (*ii*) TPS and (*iii*) TPP gene family effects, and (*iv*) combining the whole genome variation with the effects of the TPS and TPP gene families.

## Acknowledgements

Rothamsted Research receives strategic funding from the Biotechnological and Biological Sciences Research Council of the UK. We acknowledge International Wheat Yield Partnership grant (BB/S01280X/1) and Designing Future Wheat Institute Strategic Programme (BB/P016855/1). GM and MPR were supported by the International Wheat Yield Partnership (IWYP-HUB) and by the Sustainable Modernization of Traditional Agriculture (MasAgro) an initiative from the Secretariat of Agriculture and Rural Development (SADER) and CIMMYT. A.A.I received support from the Nottingham-Rothamsted Doctoral Training Partnership.

## Author contributions

D.H.L performed statistical and quantitative genetic analysis. A.W, C.A.G, and A.A.I contributed to the exome data analysis and interpretation of the biological results. G.M and M.R conducted the field experiment, assisted with the phenotypic data, and generated the 35 K SNP Chip data. R.J and A.H designed enrichment capture probe set and performed bioinformatics analysis. K.H.P assisted with the bioinformatics analysis. D.H.L and M.J.P. wrote the article, which all authors edited and approved.

## Data availability

The exome-capture data, as well as the R scripts used for most of the analyses in this study, can be found at https://github.com/DaniloLyra/exome_HiBAP_data. The genotypic (35K SNP Chip) and phenotypic data are available in Molero *et al*. (2019).

## Supporting Information

**Figure S1** Manhattan and QQ plots from the gene-based association analysis in the wheat HiBAP panel.

**Figure S2** Intragenic structure of linkage disequilibrium (LD) in trehalose family genes in the wheat HiBAP panel.

**Table S1** Results from the single variant analysis, epistasis, signature of selection, heritability of single gene and gene family, and gene-based prediction.

**Table S2** Exome-capture summary of trehalose phosphate synthase (TPS) and trehalose phosphate phosphatase (TPP) genes.

**Table S3** List of variants significantly associated with source-and sink-related traits from the single variant analysis.

**Methods S1** Partitioning the heritability per single gene.

**Methods S2** Partitioning the heritability for TPS and TPP gene family.

**Methods S3** Gene-based predictive models.

