## Supplementary Information for "Gene-based mapping of trehalose biosynthetic pathway genes reveals association with source- and sink-related yield traits in a spring wheat panel"

**Supporting Information**

**Figure S1** Manhattan and QQ plots from the gene-based association analysis in the wheat HiBAP panel.

**Figure S2** Knowledge network using trehalose phosphate synthase (TPS) and trehalose phosphate phosphatase (TPP) genes.

**Figure S3** Intragenic structure of linkage disequilibrium (LD) in trehalose family genes in the wheat HiBAP panel.

**Table S1** Results from the single variant analysis, epistasis, signature of selection, heritability of individual gene and gene family, and gene-based prediction.

**Table S2** Exome-capture summary of trehalose phosphate synthase (TPS) and trehalose phosphate phosphatase (TPP) genes.

**Table S3** List of variants significantly associated with source- and sink-related traits from the single variant analysis.

**Methods S1** Partitioning the heritability per single (local) gene.

**Methods S2** Partitioning the variance explained for TPS and TPP gene family within elite and exotic subpopulations.

**Methods S3** Gene-based predictive models.

**Supplementary figure legend**

**Figure S1.** Manhattan and QQ plots from the gene-based association analysis in the wheat HiBAP panel. The *x*-axis shows genomic position (chromosomes 1A–7B), and the *y*-axis shows statistical significance [–log10(*P*)]. Dotted line indicates significance level for Bonferroni correction (red, α=0.05) and False Discovery Rate (orange, α=0.05). Each dot represents a gene. Gene-based models are the sequence kernel association test (SKAT), optimized SKAT (SKAT-O), and multiple linear regression (MLR). Significant gene names are shown by black arrows. Gene families are trehalose phosphate synthase (TPS) and trehalose phosphate phosphatase (TPP).

**Figure S2.** Intragenic structure of linkage disequilibrium (LD) in two trehalose family genes in the wheat HiBAP panel. (a) trehalose phosphate synthase (TPS) and (b) trehalose phosphate phosphatase (TPP) gene family are shown in the panel. LD decay as squared correlations ($r^2$) of pairwise SNP LD against distance in base pairs (bp). Curves show nonlinear regression of $r^2$ on distance.

### Figure S1

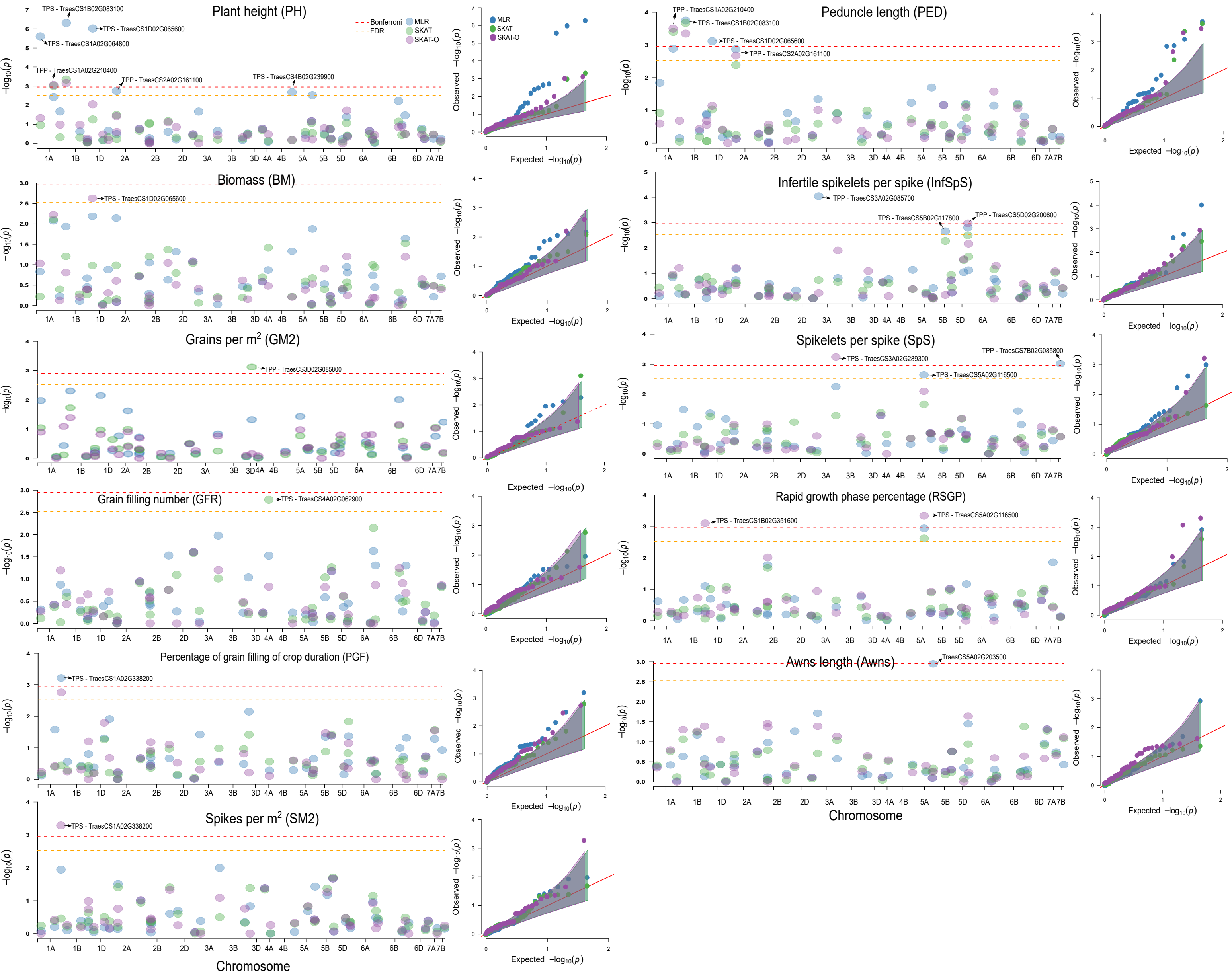

Plant height (PH)
TPS - TraesCS1B02G083100
TPS - TraesCS1D02G065600
TPS - TraesCS1A02G064800
TPP - TraesCS1A02G210400
TPP - TraesCS2A02G161100
TPS - TraesCS4B02G239900
Bonferroni
FDR
MLR
SKAT
SKAT-O
Peduncle length (PED)
TPP - TraesCS1A02G210400
TPS - TraesCS1B02G083100
TPS - TraesCS1D02G065600
TPP - TraesCS2A02G161100
Biomass (BM)
TPS - TraesCS1D02G065600
Infertile spikelets per spike (InfSpS)
TPP - TraesCS3A02G085700
TPS - TraesCS5B02G117800
TPP - TraesCS5D02G200800
Grains per m² (GM2)
TPP - TraesCS3D02G085800
Spikelets per spike (SpS)
TPP - TraesCS7B02G085800
TPS - TraesCS3A02G289300
TPS - TraesCS5A02G116500
Grain filling number (GFR)
TPS - TraesCS4A02G062900
Rapid growth phase percentage (RSGP)
TPS - TraesCS1B02G351600
TPS - TraesCS5A02G116500
Percentage of grain filling of crop duration (PGF)
TPS - TraesCS1A02G338200
Awns length (Awns)
TraesCS5A02G203500
Spikes per m² (SM2)
TPS - TraesCS1A02G338200
−log10(p)
Observed −log10(p)
Expected −log10(p)
Chromosome
1A 1B 1D 2A 2B 2D 3A 3B 3D 4A 4B 5A 5B 5D 6A 6B 6D 7A 7B

Figure S2

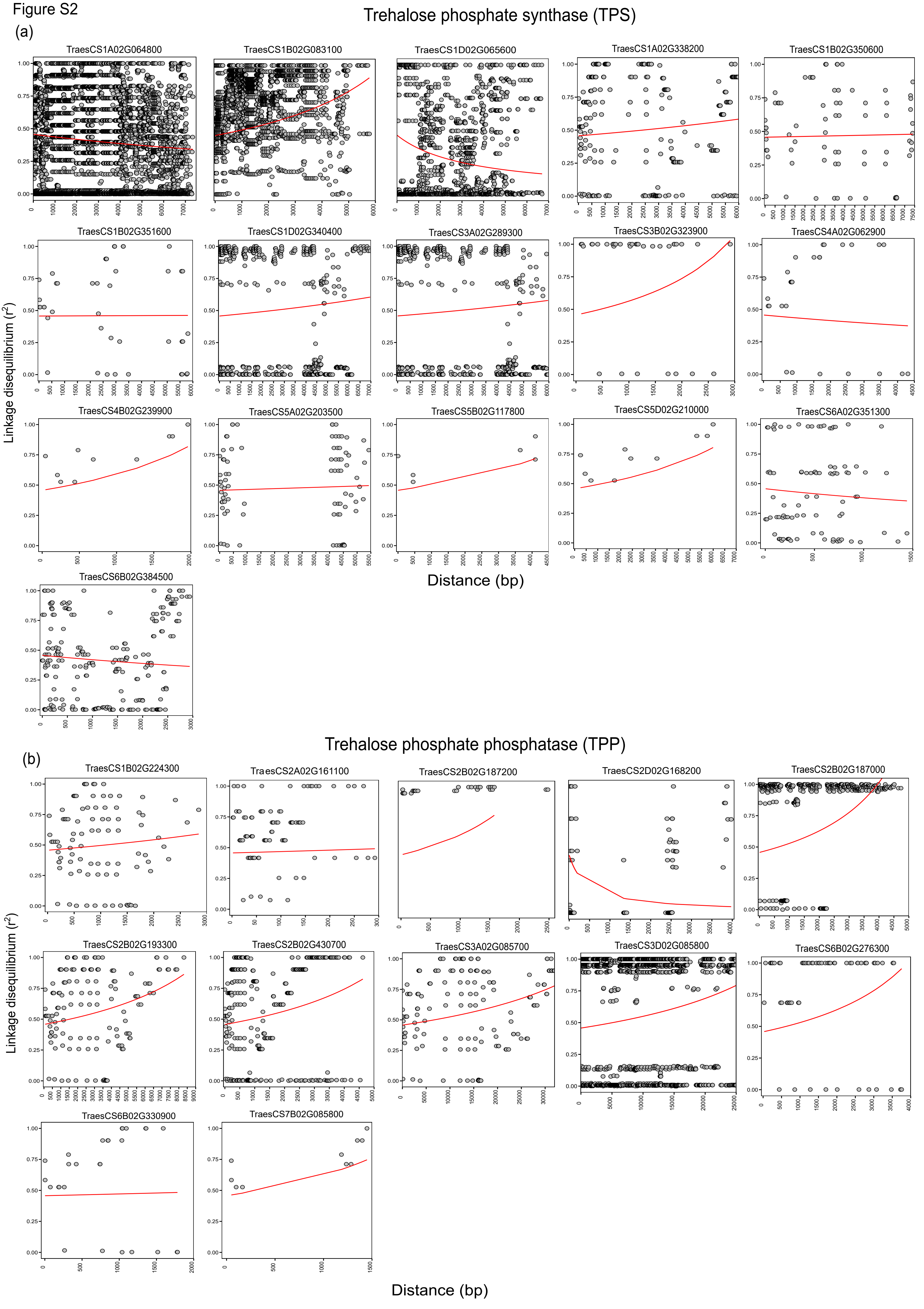
Trehalose phosphate synthase (TPS)
(a)
TraesCS1A02G064800
TraesCS1B02G083100
TraesCS1D02G065600
TraesCS1A02G338200
TraesCS1B02G350600
TraesCS1B02G351600
TraesCS1D02G340400
TraesCS3A02G289300
TraesCS3B02G323900
TraesCS4A02G062900
TraesCS4B02G239900
TraesCS5A02G203500
TraesCS5B02G117800
TraesCS5D02G210000
TraesCS6A02G351300
TraesCS6B02G384500
Linkage disequilibrium (r²)
Distance (bp)
Trehalose phosphate phosphatase (TPP)
(b)
TraesCS1B02G224300
TraesCS2A02G161100
TraesCS2B02G187200
TraesCS2D02G168200
TraesCS2B02G187000
TraesCS2B02G193300
TraesCS2B02G430700
TraesCS3A02G085700
TraesCS3D02G085800
TraesCS6B02G276300
TraesCS6B02G330900
TraesCS7B02G085800
Linkage disequilibrium (r²)
Distance (bp)

**Table S2** Exome-capture summary of trehalose phosphate synthase (TPS) and trehalose phosphate phosphatase (TPP) genes in the wheat HiBAP panel

| Gene family | Gene class | Gene ID[1] | Gene length[2] | No. variants (cleaned)[3] | No. of variants (original)[4] | $d_N$[5] | $d_S$[6] | MAF[7] | PIC[8] |
|---|---|---|---|---|---|---|---|---|---|
| Trehalose phosphate synthase | TPS1 | TraesCS1A02G064800[a] | 8.59 | 136 | 173 | 27 | 14 | 0.06±0.00 | 0.09±0.00 |
| | TPS1 | TraesCS1B02G083100[a] | 6.79 | 89 | 95 | 13 | 11 | 0.09±0.00 | 0.12±0.00 |
| | TPS1 | TraesCS1D02G065600[a] | 8.12 | 67 | 96 | 11 | 7 | 0.04±0.00 | 0.07±0.00 |
| | TPS1 | TraesCS1B02G351600 | 6.76 | 10 | 11 | 0 | 0 | 0.04±0.00 | 0.09±0.00 |
| | TPS6 | TraesCS4A02G062900[b] | 4.36 | 8 | 17 | 1 | 4 | 0.08±0.01 | 0.14±0.02 |
| | TPS6 | TraesCS4B02G239900[b] | 4.2 | 5 | 9 | 1 | 2 | 0.15±0.01 | 0.22±0.01 |
| | TPS6 | TraesCS5A02G203500[c] | 3.91 | 14 | 25 | 2 | 4 | 0.09±0.02 | 0.13±0.02 |
| | TPS6 | TraesCS5B02G202200[c] | 3.32 | 1 | 9 | 0 | 0 | 0.03±0.00 | 0.05±0.00 |
| | TPS6 | TraesCS5D02G210000[c] | 4.3 | 5 | 7 | 0 | 0 | 0.03±0.00 | 0.06±0.00 |
| | TPS7 | TraesCS1A02G338200[d] | 5.58 | 26 | 45 | 4 | 4 | 0.29±0.01 | 0.30±0.01 |
| | TPS7 | TraesCS1B02G350600[d] | 5.73 | 13 | 19 | 1 | 2 | 0.03±0.00 | 0.05±0.00 |
| | TPS7 | TraesCS1D02G340400[d] | 5.66 | 35 | 51 | 4 | 5 | 0.28±0.03 | 0.25±0.02 |
| | TPS7 | TraesCS3A02G289300[e] | 5.27 | 19 | 30 | 5 | 3 | 0.08±0.01 | 0.13±0.01 |
| | TPS7 | TraesCS3B02G323900[e] | 5.13 | 9 | 13 | 2 | 1 | 0.04±0.01 | 0.08±0.02 |
| | TPS7 | TraesCS3D02G289100[e] | 5.17 | 1 | 1 | 0 | 0 | 0.13±0.00 | 0.20±0.00 |
| | TPS7 | TraesCS5A02G116500[f] | 5.01 | 2 | 3 | 0 | 0 | 0.14±0.01 | 0.21±0.01 |
| | TPS7 | TraesCS5B02G117800[f] | 4.79 | 4 | 8 | 0 | 0 | 0.01±0.00 | 0.03±0.00 |
| | TPS7 | TraesCS5D02G129600[f] | 4.78 | 2 | 16 | 0 | 0 | 0.02±0.00 | 0.04±0.00 |
| | TPS11 | TraesCS6A02G351300[g] | 3.9 | 14 | 22 | 0 | 0 | 0.07±0.01 | 0.11±0.02 |
| | TPS11 | TraesCS6B02G384500[g] | 4.26 | 23 | 48 | 2 | 3 | 0.05±0.01 | 0.09±0.01 |
| | TPS11 | TraesCS6D02G334000[g] | 3.99 | - | 3 | 0 | 0 | - | - |
| Trehalose phosphate phosphatase | - | TraesCS1A02G210400[a] | 4.18 | 4 | 8 | 0 | 0 | 0.05±0.00 | 0.09±0.00 |
| | - | TraesCS1B02G224300[a] | 2.86 | 14 | 15 | 0 | 0 | 0.18±0.03 | 0.22±0.02 |
| | - | TraesCS1D02G213700[a] | 4.02 | 3 | 6 | 0 | 0 | 0.08±0.00 | 0.13±0.00 |
| | - | TraesCS2A02G161100[b] | 3.04 | 14 | 21 | 2 | 5 | 0.21±0.02 | 0.26±0.02 |
| | - | TraesCS2B02G187200[b] | 3.33 | 8 | 13 | 0 | 0 | 0.05±0.04 | 0.06±0.02 |
| | - | TraesCS2D02G168200[b] | 3.27 | 15 | 18 | 0 | 0 | 0.10±0.04 | 0.10±0.03 |
| | - | TraesCS2A02G161000[c] | 3.32 | - | 1 | 0 | 0 | - | - |
| | - | TraesCS2B02G187000[c] | 3.44 | 33 | 36 | 2 | 1 | 0.16±0.01 | 0.20±0.02 |
| | - | TraesCS2D02G168300[c] | 3.52 | - | 1 | 0 | 0 | - | - |
| | - | TraesCS2A02G167100[d] | 6.62 | 3 | 5 | 0 | 0 | 0.24±0.00 | 0.3±0.00 |
| | - | TraesCS2B02G193300[d] | 21.7 | 18 | 51 | 0 | 0 | 0.06±0.01 | 0.09±0.01 |
| | - | TraesCS2A02G412100[e] | 3.52 | 1 | 1 | 0 | 0 | 0.23±0.00 | 0.29±0.02 |
| | - | TraesCS2B02G430700[e] | 3.32 | 28 | 29 | 3 | 1 | 0.09±0.00 | 0.14±0.01 |
| | - | TraesCS2D02G409300[e] | 3.26 | 1 | 2 | 0 | 0 | 0.45±0.00 | 0.37±0.00 |
| | - | TraesCS3A02G085700[f] | 32.3 | 19 | 66 | 0 | 0 | 0.10±0.03 | 0.11±0.03 |
| | - | TraesCS3D02G085800[f] | 24.2 | 65 | 144 | 6 | 4 | 0.06±0.00 | 0.10±0.01 |
| | - | TraesCS5A02G190000[g] | 2.32 | 2 | 3 | 0 | 0 | 0.14±0.02 | 0.21±0.02 |
| | - | TraesCS5B02G193100[g] | 2.3 | 1 | 2 | 0 | 0 | 0.02±0.00 | 0.04±0.00 |
| | - | TraesCS5D02G200800[g] | 2.39 | 2 | 4 | 0 | 0 | 0.17±0.01 | 0.20±0.13 |
| | - | TraesCS6A02G248400[h] | 3.76 | 2 | 14 | 0 | 0 | 0.03±0.00 | 0.06±0.00 |
| | - | TraesCS6B02G276300[h] | 4.27 | 12 | 23 | 0 | 0 | 0.06±0.02 | 0.10±0.02 |
| | - | TraesCS6D02G230500[h] | 3.92 | 2 | 2 | 0 | 0 | 0.03±0.02 | 0.06±0.04 |
| | - | TraesCS6A02G301800[i] | 2.95 | 3 | 25 | 0 | 0 | 0.07±0.04 | 0.12±0.05 |
| | - | TraesCS6B02G330900[i] | 3.25 | 8 | 13 | 0 | 0 | 0.06±0.02 | 0.10±0.03 |
| | - | TraesCS6D02G281100[i] | 2.96 | 1 | 2 | 0 | 0 | 0.06±0.00 | 0.10±0.00 |
| | - | TraesCS7A02G180800[j] | 2.57 | 2 | 7 | 3 | 1 | 0.07±0.01 | 0.12±0.03 |
| | - | TraesCS7B02G085800[j] | 2.61 | 5 | 10 | 0 | 0 | 0.06±0.02 | 0.09±0.03 |

[1]Wheat gene ID at EnsemblPlants (IWGSC RefSeq v1.0 annotation). [a-j] Same letter indicates homoelogues genes

[2]Gene length in kilobase (Kb)

[3]Number of variants inside the gene after applying for quality control (minor allele frequency <1% and call rate <95%)

[4]Number of variants inside the gene without applying quality control

[5-6]Total number of nonsynonymous ($d_N$) and synonymous ($d_S$) substitutions in the original data set using the EnsemblPlants Variant Effect Predictor (VEP) tool

[7]Minor allele frequency (MAF). Values are mean±standard errors from all variants inside the gene

[8]Polymorphism information content (PIC). Values are mean±standard errors from all variants inside the gene

**Table S3** List of variants significantly associated with source- and sink-related traits from the single variant analysis in the wheat HiBAP panel

| Trait[1] | Variant ID[2] | Gene class[3] | Gene ID[4] | MAF[5] | *P* value[6] | Beta[7] | Annotation[8] | SIFT[9] |
|---|---|---|---|---|---|---|---|---|
| PH | chr1D-47313394[a] | *TPS1* | TraesCS1D02G065600 | 0.01 | $4.25\times10^{-07}$ | 21.8 | Missense | Deleterious |
| | chr1D-47313412[a] | *TPS1* | TraesCS1D02G065600 | 0.01 | $4.25\times10^{-07}$ | 21.8 | Missense | Tolerated |
| | chr1B-67021268[a] | *TPS1* | TraesCS1B02G083100 | 0.02 | $4.88\times10^{-07}$ | 10.4 | Intron | - |
| | chr1A-47193551[b] | *TPS1* | TraesCS1A02G064800 | 0.01 | $2.42\times10^{-04}$ | 12.1 | Intron | - |
| PED | chr1D-47313394[a] | *TPS1* | TraesCS1D02G065600 | 0.01 | $6.81\times10^{-06}$ | 10.7 | Missense | Deleterious |
| | chr1D-47313412[a] | *TPS1* | TraesCS1D02G065600 | 0.01 | $6.81\times10^{-06}$ | 10.7 | Missense | Tolerated |
| | chr1B-67021268[a] | *TPS1* | TraesCS1B02G083100 | 0.02 | $1.77\times10^{-05}$ | 4.87 | Intron | - |
| GM2 | chr1A-47193551[a] | *TPS1* | TraesCS1A02G064800 | 0.01 | $6.05\times10^{-05}$ | -3395 | Intron | - |
| | chr1D-47313394[b] | *TPS1* | TraesCS1D02G065600 | 0.02 | $1.81\times10^{-04}$ | -3967 | Missense | Deleterious |
| | chr1D-47313412[b] | *TPS1* | TraesCS1D02G065600 | 0.02 | $1.81\times10^{-04}$ | -3967 | Missense | Tolerated |
| GFR | chr6B-659234423[a] | *TPS11* | TraesCS6B02G384500 | 0.01 | $3.56\times10^{-05}$ | -2.63 | Intron | - |
| | chr1A-527948409[b] | *TPS7* | TraesCS1A02G338200 | 0.01 | $1.62\times10^{-04}$ | -3.24 | Upstream | - |
| | chr1A-527949030[b] | *TPS7* | TraesCS1A02G338200 | 0.01 | $1.62\times10^{-04}$ | -3.24 | Upstream | - |
| | chr1A-527955187[b] | *TPS7* | TraesCS1A02G338200 | 0.01 | $1.62\times10^{-04}$ | -3.24 | Synonymous | - |
| PGF | chr1A-527948409[a] | *TPS7* | TraesCS1A02G338200 | 0.01 | $2.59\times10^{-05}$ | 4.52 | Upstream | - |
| | chr1A-527949030[a] | *TPS7* | TraesCS1A02G338200 | 0.01 | $2.59\times10^{-05}$ | 4.52 | Upstream | - |
| | chr1A-527955187[a] | *TPS7* | TraesCS1A02G338200 | 0.01 | $2.59\times10^{-05}$ | 4.52 | Synonymous | - |
| SM2 | chr1A-527948409[a] | *TPS7* | TraesCS1A02G338200 | 0.01 | $5.26\times10^{-06}$ | 93.8 | Upstream | - |
| | chr1A-527949030[a] | *TPS7* | TraesCS1A02G338200 | 0.01 | $5.26\times10^{-06}$ | 93.8 | Upstream | - |
| | chr1A-527955187[a] | *TPS7* | TraesCS1A02G338200 | 0.01 | $5.26\times10^{-06}$ | 93.8 | Synonymous | - |
| | chr3A-517661865[b] | *TPS7* | TraesCS3A02G289300 | 0.01 | $1.22\times10^{-04}$ | 52.4 | 3' UTR | - |
| GWSP | chr1A-527948409[b] | *TPS7* | TraesCS1A02G338200 | 0.01 | $1.62\times10^{-04}$ | -0.56 | Upstream | - |
| | chr1A-527949030[b] | *TPS7* | TraesCS1A02G338200 | 0.01 | $1.62\times10^{-04}$ | -0.56 | Upstream | - |
| | chr1A-527955187[b] | *TPS7* | TraesCS1A02G338200 | 0.01 | $1.62\times10^{-04}$ | -0.56 | Synonymous | - |

[1]Traits are plant height (PH, cm), peduncle length (PED, cm), grains per $m^2$ (GM2), grain filling rate (GFR, yield/grain filling duration, $g/m^2$/day), percentage of grain filling of crop duration (PGF), spikes per $m^2$ (SM2), and grain weight per spike (GWSP, g)

[2]Chromosome and position (bp) of each variant. Significant variants from [a]Bonferroni and [b]FDR correction (α=0.05)

[3]Trehalose phosphate synthase (TPS) and trehalose phosphate phosphatase (TPP) gene family class

[4]Wheat gene description at EnsemblPlants (IWGSC RefSeq v1.0 annotation)

[5]Minor allele frequency (MAF)

[6]*P*-value from the linear mixed model (MLM)

[7]Allelic substitution effect (beta coefficient)

[8]Annotation using the EnsemblPlants Variant Effect Predictor (VEP) tool

[9]Tolerated and deleterious missense mutations in a gene from the Sorting Intolerant From Tolerant (SIFT) score

**Methods S1**. Partitioning the heritability per single gene

We combined two kernels into the mixed linear model (MLM) to estimate the proportion of the phenotypic variance explained (i.e. genomic heritability) per gene. Similar method has often been called Regional Heritability Mapping (RHM) (Nagamine *et al.*, 2012; Uemoto *et al.*, 2013). We used the variants per gene to build a local genomic relationship matrix (GRM) and combined with the effects of the global GRM from the genome-wide markers (35K SNP Chip).

We fitted the following mixed linear model:

$$\hat{\boldsymbol{y}} = \boldsymbol{X\beta} + \boldsymbol{Z_g g} + \boldsymbol{Z_l l} + \boldsymbol{\varepsilon}, \quad (1)$$

where $\hat{\boldsymbol{y}}$ was a vector of phenotypic values, $\boldsymbol{\beta}$ was the vector of fixed effects (with and without PC adjustment), $\boldsymbol{g}$ was the vector of random additive genetic effects of the global genome-wide SNP markers, $\boldsymbol{l}$ was the vector of random additive genetic effects of the local gene region markers, and $\boldsymbol{\varepsilon}$ was a vector of random residuals. The incidence matrices for $\boldsymbol{\beta}$, $\boldsymbol{g}$, and $\boldsymbol{l}$ were $\boldsymbol{X}$, $\boldsymbol{Z_g}$, and $\boldsymbol{Z_l}$, respectively. The distributions of random effects were assumed to be $\boldsymbol{g} \sim N(\boldsymbol{0}, \sigma_g^2 \boldsymbol{G_g})$, $\boldsymbol{l} \sim N(\boldsymbol{0}, \sigma_l^2 \boldsymbol{G_l})$, and $\boldsymbol{\varepsilon} \sim N(\boldsymbol{0}, \sigma_\varepsilon^2 \boldsymbol{I})$, where $\boldsymbol{I}$ was the identity matrix, $\boldsymbol{G_g}$ was the global additive GRM calculated using $\boldsymbol{G_g} = \boldsymbol{WW'}/g$, and $\boldsymbol{G_l}$ is the local additive GRM calculated the same way as $\boldsymbol{G_g}$. $\boldsymbol{W}$ is a $n \times g$ matrix of scaled and centered markers from $n$ individuals and $g$ is the total number of markers. To build the $\boldsymbol{W}$ matrix we used a genotypic incidence matrix coded as 2 for homozygote $A_1A_1$, 1 for heterozygote $A_1A_2$, and 0 for homozygote $A_2A_2$. We extracted genomic estimates of the additive global whole genomic ($\sigma_g^2$), local gene ($\sigma_l^2$), and residual ($\sigma_\varepsilon^2$) variances, enabling the calculation of local gene heritability as $h_l^2 = \sigma_l^2/(\sigma_g^2 + \sigma_l^2 + \sigma_\varepsilon^2)$ and global whole genomic heritability as $h_g^2 = \sigma_g^2/(\sigma_g^2 + \sigma_l^2 + \sigma_\varepsilon^2)$. We tested the presence of regional/local gene variance ($\sigma_l^2$) using a likelihood ratio test $[LRT = -2\ln(L_0/L_1)]$, where $L_0$ and $L_1$ are the likelihood values for the hypothesis of absence ($H_0: \sigma_l^2 = 0$ ) or presence ($H_1: \sigma_l^2 > 0$) of regional variance, respectively. We adjusted the $P$-values for multiple comparisons to control for type I error at $\alpha = 0.05$ using the Bonferroni procedure (the number of genes tested was considered to set the threshold).

**Methods S2**. Partitioning the heritability of TPS and TPP gene family

We combined three kernels into the MLM to estimate the proportion of the phenotypic variance explained of TPS and TPP gene families. We used the variants of each gene family (TPS and TPP) to build a local GRM and combined with the effects of the global GRM from the genome-wide markers.

We fitted the following mixed linear model:

$$\boldsymbol{\hat{y}} = \boldsymbol{X\beta} + \boldsymbol{Z_g g} + \boldsymbol{Z_{TPS} TPS} + \boldsymbol{Z_{TPP} TPP} + \boldsymbol{\varepsilon}, \quad (2)$$

where $\boldsymbol{\hat{y}}$ was a vector of phenotypic values, $\boldsymbol{\beta}$ was the vector of fixed effects, $\boldsymbol{g}$ was the vector of random additive genetic effects of the global genome-wide SNP markers, $\boldsymbol{TPS}$ and $\boldsymbol{TPP}$ were the vectors of random additive genetic effects of the local TPS and TPP gene family markers, and $\boldsymbol{\varepsilon}$ was a vector of random residuals. The incidence matrices for $\boldsymbol{\beta}$, $\boldsymbol{g}$, $\boldsymbol{TPS}$, and $\boldsymbol{TPP}$ were $\boldsymbol{X}$, $\boldsymbol{Z_g}$, $\boldsymbol{Z_{TPS}}$, and $\boldsymbol{Z_{TPP}}$, respectively. The distributions of random effects were assumed to be $\boldsymbol{g} \sim N(\boldsymbol{0}, \sigma_g^2 \boldsymbol{G_g})$, $\boldsymbol{TPS} \sim N(\boldsymbol{0}, \sigma_{TPS}^2 \boldsymbol{G_{TPS}})$, $\boldsymbol{TPP} \sim N(\boldsymbol{0}, \sigma_{TPP}^2 \boldsymbol{G_{TPP}})$, and $\boldsymbol{\varepsilon} \sim N(\boldsymbol{0}, \sigma_\varepsilon^2 \boldsymbol{I})$, where $\boldsymbol{I}$ was the identity matrix, $\boldsymbol{G_g}$ was the global additive GRM calculated using $\boldsymbol{G_g} = \boldsymbol{WW'}/g$, and $\boldsymbol{G_{TPS}}$ and $\boldsymbol{G_{TPP}}$ are the local additive GRM calculated the same way as $\boldsymbol{G_g}$. $\boldsymbol{W}$ is a $n \times g$ matrix of scaled and centered markers from $n$ individuals and $g$ is the total number of markers. To build the $\boldsymbol{W}$ matrix we used a genotypic incidence matrix coded as 2 for homozygote $A_1A_1$, 1 for heterozygote $A_1A_2$, and 0 for homozygote $A_2A_2$. We extracted genomic estimates of the additive global whole genomic ($\sigma_g^2$), local TPS and TPP gene family ($\sigma_{TPS}^2$ and $\sigma_{TPP}^2$), and residual ($\sigma_\varepsilon^2$) variances, enabling the calculation of local TPS gene family heritability as $h_{TPS}^2 = \sigma_{TPS}^2/(\sigma_g^2 + \sigma_{TPS}^2 + \sigma_{TPP}^2 + \sigma_\varepsilon^2)$, local TPP gene family heritability as $h_{TPP}^2 = \sigma_{TPP}^2/(\sigma_g^2 + \sigma_{TPS}^2 + \sigma_{TPP}^2 + \sigma_\varepsilon^2)$, and global whole genomic heritability as $h_g^2 = \sigma_g^2/(\sigma_g^2 + \sigma_{TPS}^2 + \sigma_{TPP}^2 + \sigma_\varepsilon^2)$.

**Methods S3**. Gene-based predictive models

We predicted the phenotypic traits using the trehalose biosynthetic pathway genes (TPS and TPP kernels) individually and jointly with the whole genome markers (35K SNP Chip). We used population structure variables (matrix of zeros and ones based on Molero *et al.* (2019) group clustering) as fixed covariates in the model (Lyra *et al.*, 2018). Predictive ability (*r*) was calculated as the Pearson correlation between adjusted values and genomic estimated breeding values in 50 replications from independent validation scenarios (Albrecht *et al.*, 2014), randomly sampling 75% of the genotypes (*n*=110) to form a training set, while the remaining 25% (*n*=37) were used as a validation set. We applied Fisher's Z transformation of the predictive abilities and compared them among models using Tukey's test at α=0.05. All prediction analyses were performed using the BGLR R package (Perez & de los Campos, 2014), using 60 000 Markov Chain Monte Carlo (MCMC) iterations, with 15 000 iterations for burn-in, and keeping only one from every five consecutive iterations to minimize auto-correlation.

First, we used the additive genome-wide marker effects as predictors by fitting the following GBLUP model:

$$\boldsymbol{\hat{y}} = \boldsymbol{X\beta} + \boldsymbol{Z_g g} + \boldsymbol{\varepsilon}, \qquad (3)$$

where $\boldsymbol{\hat{y}}$, $\boldsymbol{\beta}$, $\boldsymbol{g}$, and $\boldsymbol{\varepsilon}$ are the same as those defined in the Eq. (1).

Second, we used the local additive TPS gene family effects as predictors by fitting the following GBLUP model:

$$\boldsymbol{\hat{y}} = \boldsymbol{X\beta} + \boldsymbol{Z_{TPS} TPS} + \boldsymbol{\varepsilon}, \qquad (4)$$

where $\boldsymbol{\hat{y}}$, $\boldsymbol{\beta}$, $\boldsymbol{TPS}$, and $\boldsymbol{\varepsilon}$ are the same as those defined in the Eq. (2). The single kernel TPP gene family was used the same way as Eq. (4).

Finally, we combine all kernels (global whole genomic, TPS, and TPP genic effects) in the GBLUP model using the same model as Eq. (2).

**Observation.** For the Methods S1-S3, we used the phenotypic traits of 147 individuals (only these genotypes were available in the 35K SNP Chip ID). Also, genes with only one variant were removed from the analyses. For the Methods S1 and S2, we used the complete set of individuals (149 lines), and elite and exotic subpopulations independently.
